# C/EBPδ demonstrates a dichotomous role in tumor initiation and promotion of epithelial carcinoma

**DOI:** 10.1101/528885

**Authors:** Ramlogan Sowamber, Rania Chehade, Mahmoud Bitar, Leah Dodds, Anca Milea, Brian Slomovitz, Patricia A Shaw, Sophia HL George

## Abstract

*C/EBPδ* (CEBPD), a gene part of the highly conserved basic-leucine zipper (b-ZIP) domain of transcriptional factors, is downregulated in 65% of high grade serous carcinoma of the ovary (HGSC). Overexpression of *C/EBPδ* in different tumors as glioblastoma and breast cancer either promotes tumor progression or inhibits growth. Despite these contradictory roles in different cancer types, we show that *C/EBPδ* overexpression has a consistent function of downregulating proliferation and promoting migration in fallopian tube epithelial cells (FTE). We show that the FTE have both mesenchymal and epithelial cell characteristics. Further, our data supports a role for *C/EBPδ* as an early regulatory transcriptional factor that promotes a mesenchymal to epithelial (MET) phenotype by upregulating E-cadherin and downregulating vimentin and N-cadherin in FTE cells. We demonstrate that overexpression of *C/EBPδ* in ovarian and breast cancer cell lines have consistent effects and phenotype as the FTE cells. Our findings suggest a role for *C/EBPδ* in the early events of ovarian serous carcinogenesis which may be used to help further understand how the disease develops from a premalignant cells.

## Introduction

High-grade serous ovarian cancer (HGSC) remains the most fatal gynecological malignancy and accounts for the majority of deaths due to ovarian cancer [1]. Improving early detection, prevention and overall prognosis was, limited by a lack of understanding of the etiology of HGSC. Now, significant data suggests the distal end of the fallopian tube is the site of origin for HGSC [2]. Detailed histopathological examination of the fallopian tube epithelium in *BRCA* mutation carriers undergoing prophylactic bilateral salpingo-oophorectomy led to the identification of precursor lesions [3, 4].

A pre-neoplastic lesion called the p53 signature is the earliest mutational and genomic event described in the gradual steps of HGSC development [5]. The acquisition of somatic *TP53* mutations is followed by additional genomic alterations, cellular tufting and loss of polarity, resulting in the development of neoplastic serous tubal intraepithelial carcinoma (STIC) [6-12]. STIC lesions share multiple genomic copy number alterations, along with mutations in tumor suppressors and oncogenes, including *TP53*, *BRCA1* and *BRCA2*, *RB1*, *STK11*, *FOXO3a*, *CCNE1*, *STATHMIN1* and *hTERT*, which are observed prior to HGSC metastasis to the ovaries and peritoneal spread [6, 10, 13-19]. In this model, cells that exfoliate from the fallopian tube into the peritoneal cavity must evade anoikis and detachment associated apoptosis before attaching to the mesothelial ovarian surface [20, 21].

The fallopian tube epithelia (FTE) undergo monthly cycles of hormonally driven proliferation and differentiation. In a previous study, we identified CCAAT/enhancer binding protein delta (*C/EBPδ*), to be transcriptionally upregulated in the FTE of *BRCA1* mutation carriers and in the post-ovulatory (luteal) phase of the ovarian cycle a process linked to cytotoxic stress [22]. *C/EBPδ* is located on chromosome 8q11.21 and belongs to the superfamily of highly conserved basic-leucine zipper (b-ZIP) domain transcriptional factors [23]. It has multiple functions related to inflammation, cell cycle regulation, differentiation and metabolism [22-25]. Although C/EBPδ overexpression promotes glioblastoma progression and is associated with poor progression in pancreatic and urothelial cancers [26, 27], its overexpression in breast, prostate and myeloid cancers, inhibits growth and promotes differentiation [28-30]. Furthermore, low expression of C/EBPδ was reported in cervical, hepatocellular carcinoma [31, 32], breast [33], prostate cancer [34], and leukemia [28]. *C/EBPδ*’s pro-oncogenic/tumor suppressive function is cell type and context dependent [26, 31, 35-37]. Little is known about the role of *C/EBPδ* in the development of HGSC. The objectives of this study were to explore expression of *C/EBPδ* in HGSC tumors as well as precursor lesions and determine the effects of *C/EBPδ* on FTE cancer cell growth and migration.

## Results

### C/EBPδ is differentially expressed across serous ovarian cancer histotypes

We previously reported higher expression levels of C/EBPδ at the mRNA and protein levels in the luteal phase of the normal fallopian tube epithelia [22]. To investigate C/EBPδ protein expression levels across serous ovarian cancer histotypes, immunohistochemistry was performed on a cohort of 366 high grade serous carcinoma (HGSC) and 26 low-grade serous carcinoma (LGSC) on 5 independent tissue microarrays (TMAs). Each core was annotated to include epithelia and exclude stroma. Mean intensity and percent positivity using a nuclear stain algorithm was reported based on the following histoscores: no expression (0); low (+1); medium (+2) and high (+3) (Figure 1A). In FTE, C/EBPδ is expressed in the nuclei of secretory and ciliated cells. Seventy-six per cent (76.5%, 280/366) of HGSC cases had low or attenuated (histoscore of 0/+1) C/EBPδ protein expression whereas 24% (86/366) had a medium to high expression (histoscore of + 2/+3). Seventy-six per cent (76.9%, 20/26) of LGSC cases had medium to high expression compared to 23% (6/26) with low C/EBPδ expression. Overall, HGSC had 2-fold lower C/EBPδ protein expression relative to LGSC (p = 0.0004) (Figure 1B).

**Figure 1.**
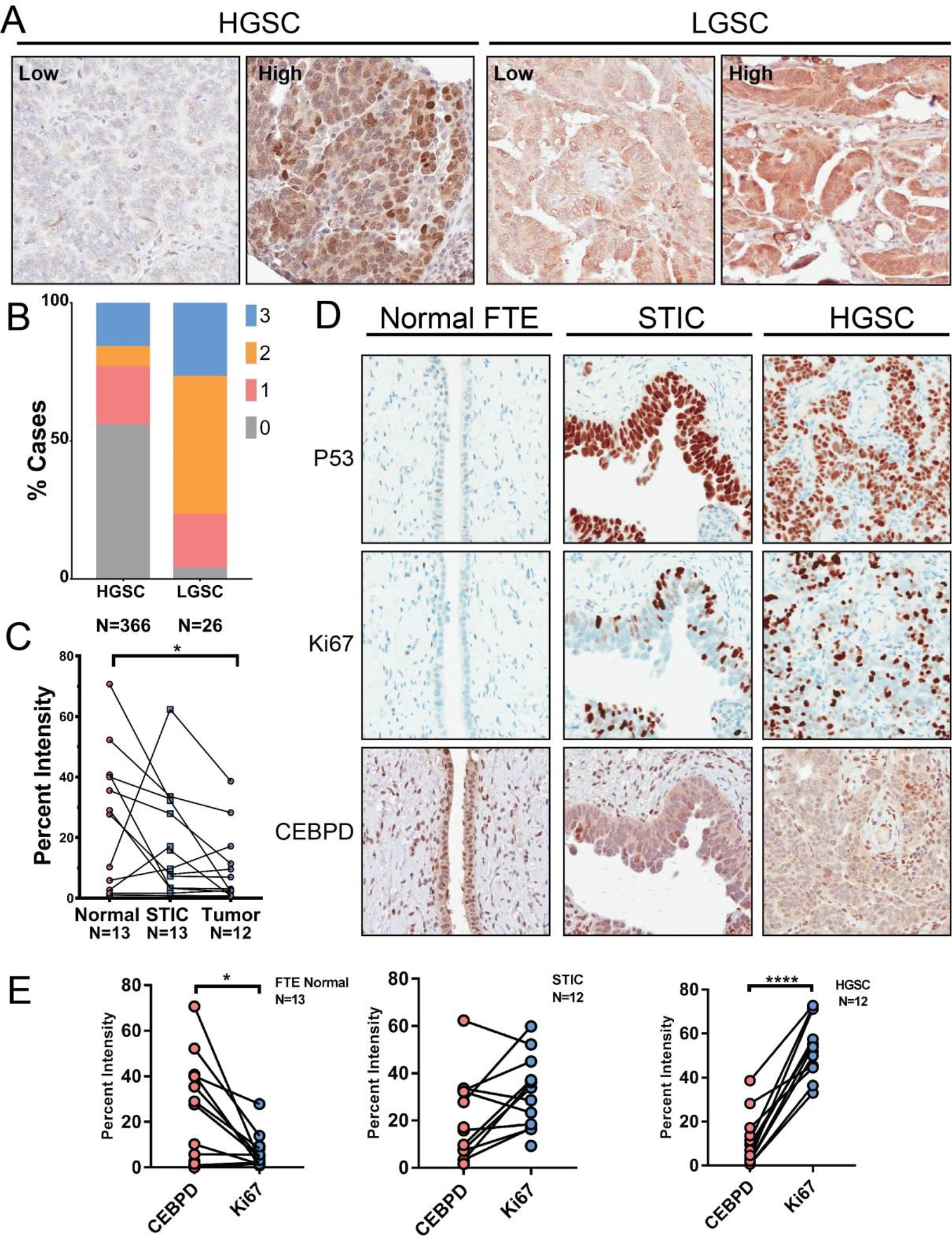
C/EBPδ is differentially expressed between serous histotypes. **A.** Representative IHC staining of C/EBPδ showing examples of HGSC and LGSC expressing low and high C/EBPδ expression, respectively. Percentage of C/EBPδ staining was scored from 0 (low) to 3 (high). **B.** Histogram shows C/EBPδ expression levels stratified by scores across a cohort of HGSC (N = 366) and LGSC (N = 26). **C.** Dot plot of representative samples shows decreasing C/EBPδ IHC staining across normal FTE (N = 13), STIC (N = 13) and HGSC (N = 12). C/EBPδ expression differs significantly between normal FTE and HGSC tumor samples. **D.** Representative IHC staining of p53, Ki67 and C/EBPδ in histologically normal fallopian tube epithelia, STIC and HGSC, respectively. **E.** Quantification of both C/EBPδ and Ki67 levels in FTE (N = 13), STIC (N = 12), and HGSC cases (N = 12) demonstrated an inverse relationship between C/EBPδ and Ki67 in normal (p = 0.01) and HGSC cases (p = 0.0001). Data are represented as mean +/- SEM and p-values are calculated using student t-tests, two-tailed (*p < 0.05; **** p < 0.0001).

To determine if C/EBPδ protein expression seen in HGSC is correlated to genomic changes, we used a cohort of HGSC genomic data (Affymetrix 6.0 SNP array) of chemotherapy naïve and stage III/IV cases (n = 75) [6, 13]. Seventeen per cent (13/75) of cases (p < 0.0001) had an amplification of the gene whereas only 2% (2/75) cases revealed a deletion; most cases (59/75) remained diploid at that genomic locus (Supplementary Figure 1A). In the TCGA ovarian cancer dataset, [16], 2.5% (8/316) of HGSC cases had an amplification and 3.1% (10/316) had a shallow deletion of *C/EBPδ* with few cases (less than 10%) having the mRNA downregulated (http://bit.ly/2DW9QPa) (Supplementary Figure 1A). Patients with high (+2/+3) C/EBPδ protein expression had no significant progression free survival (PFS, Log rank test, p = 0.319) or overall survival (OS) (Log rank test, p = 0.330) compared to low (0/+1) C/EBPδ protein expression levels (Supplementary Figure 1B).

### Reduced C/EBPδ expression levels in STIC lesions reflect early changes observed in HGSC

Since C/EBPδ levels are higher in normal FTE and differentially expressed in HGSC, we assessed C/EBPδ expression in a small cohort of matched normal FTE, STIC and HGSC (n = 13). Our data revealed that 53.8% (7/13) of normal fallopian tube cases had higher C/EBPδ expression levels (+2/+3 expression) compared to other STIC’s and HGSC, whereas in 46% (6/13) of cases, C/EBPδ expression was maintained between FTE and STIC. In normal FTE, high C/EBPδ expression was associated with low proliferation as determined by Ki67 expression (p = 0.02). In STIC, however, C/EBPδ expression was slightly lower than in normal FTE, while Ki67 expression increased relative to normal FTE (p = 0.08) and HGSCs. There were fewer C/EBPδ expressing cells in HGSC relative to highly proliferative (Ki67^+^) and p53 expressing cells (p < 0.0001) (Figure 1C-E). Overall, C/EBPδ expression was inversely related to proliferation during the transition from normal FTE, to STIC and subsequently HGSC. Given the results and the known role of C/EBPδ in cell cycle control, we sought to determine whether overexpression of C/EBPδ would regulate FTE and cancer cell growth.

### Overexpression of C/EBPδ inhibits proliferation in premalignant fallopian tube epithelia and cancer cells

#### Fallopian Tube Epithelia

To model p53 signatures *in vitro*, FTE cell lines with a p53-R175H dominant negative mutation (p53DN) and human telomerase (hTERT) over-expression were generated [6](Figure 2A). As expected from immunohistochemical data, proliferating p53DN FTE cell lines had low levels of endogenous C/EBPδ protein (Figure 2B-C). A C/EBPδ gain-of-function model was generated with a lentivirus based *pCDH-CMV-GFP-PURO* expression vector, to examine its effects on proliferation of FTE cells. Three independent FTE cell lines were transfected with either empty vector (FTE-ctrl) or C/EBPδ cDNA vector (FTE-CEBPD-OE) to generate stable lines (Figure 2A-B, Supplementary Figure 2B). C/EBPδ overexpression was confirmed by immunofluorescence (localized to the nucleus) and by western blot analyses (Figure 2B-C). To examine the role of C/EBPδ in cell cycle regulation, cell cycle analysis and growth kinetics were performed on FTE-ctrl and FTE-CEBPD-OE cell lines. Over-expression of C/EBPδ reduced cell growth compared to controls, as measured by population doubling (PD) over time (FTE19-CEBPD-OE versus control, 7.1-fold, p = 0.04; FTE57-CEBPD-OE vs control, 14.9-fold, p = 0.01; each in triplicate) (Figure 2D, Supplementary Figure 2C). Cell cycle analysis by BrdU /propidium iodide incorporation and flow cytometry in FTE cells demonstrated a propensity for C/EBPδ overexpressing cells to accumulate in the G1 phase of the cell cycle compared to FTE-ctrl cells (p = 0.0019) (Figure 2E). C/EBPδ is known to interact with *CCND1* and promotes STAT3 induced G0 cell cycle arrest [38, 39]. In addition, overexpression of C/EBPδ decreased anchorage independent growth of FTE cells compared to control cells (FTE19-p53DN-hTERT, 1.97-fold, p = 0.0009; FTE57-p53DN-hTERT, 1.41-fold, p = 0.07) (Figure 2F), but increased colony formation in one cell line (Supplementary Figure 2D). Overall, the results are consistent with decreased growth rates in 2D and 3D assays. The data show that C/EBPδ overexpression is sufficient to inhibit proliferation in FTE cells likely at the G1/S phase of the cell cycle.

**Figure 2.**
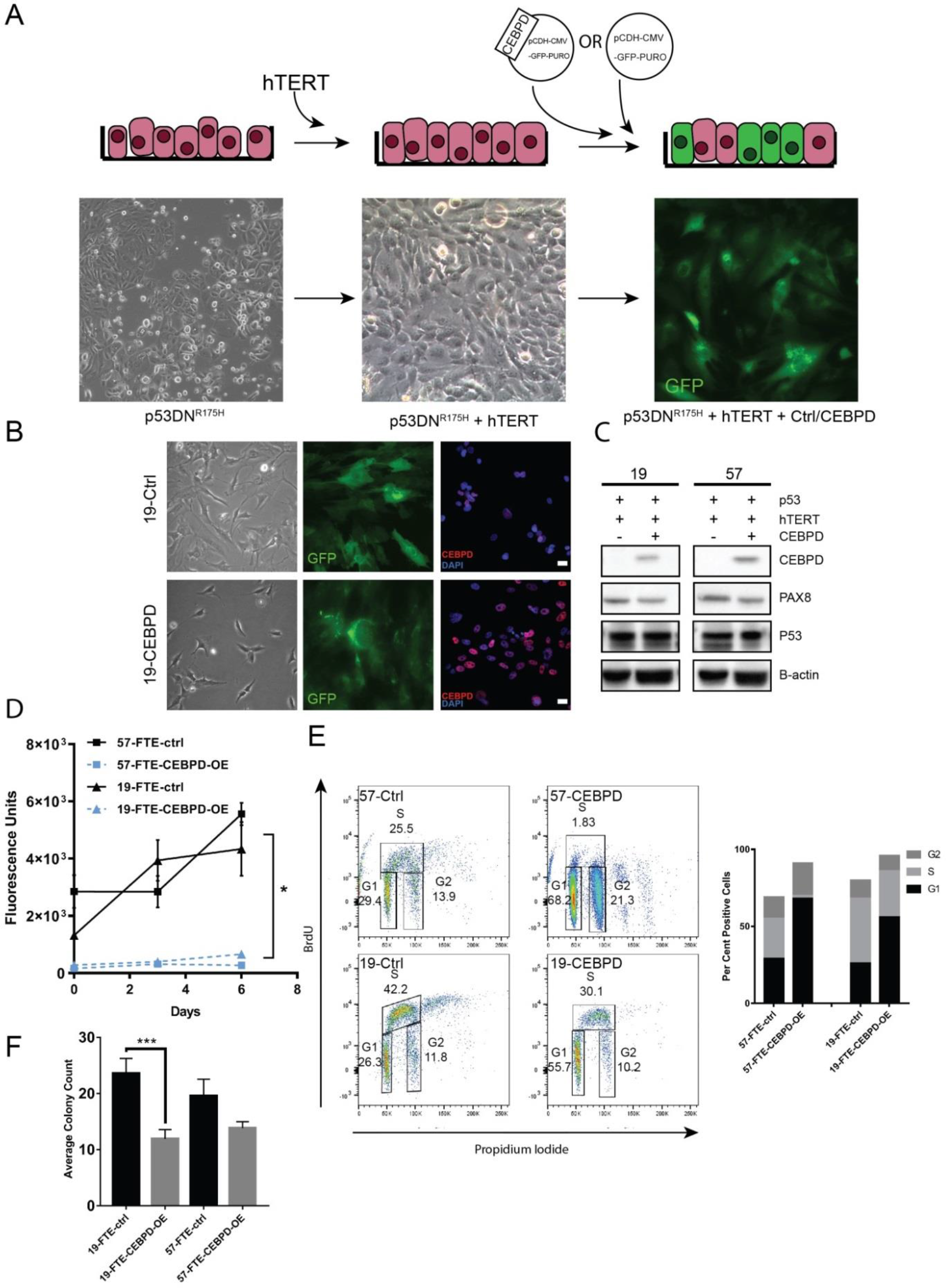
Overexpression of C/EBPδ inhibits proliferation. **A.** *In vitro* model of C/EBPδ overexpression in premalignant fallopian tube epithelia. Fallopian tube epithelial cells have a limited life span *in-vitro.* To generate premalignant cells, FTE were infected with lentivirus containing P53 dominant negative mutation (P53 DN^R175H^). Cells with P53 DN^R175H^ were selected and subsequently immortalized using htert. FTE-P53DN-htert cells were then infected with either control vector or C/EBPδ. **B.** Phase contrast microscopy images of a FTE cell line, showing control cells (19-FTE-P53DN-hTert-ctrl) and C/EBPδ overxpressing cells (19-FTE-P53DN-hTert-CEBPD-OE) with corresponding images of GFP expression (green) as a marker of successfully infected cells, and C/EBPδ expression (red) in the nucleus (blue). **C.** Confirmation of C/EBPδ protein expression by western blot in two FTE cell lines, 19 and 57. Pax 8 is a fallopian tube epithelial marker. **D.** Cell growth measured as population doublings across days in two cell lines, 19 and 57. C/EBPδ overexpression significantly reduced growth in premalignant FTE. **E.** To determine cell cycle distribution, BrdU incorporation and DNA content were analyzed by flow cytometry. Representative plot showing percentage distribution of cells across cell cycle control verses C/EBPδ overexpressing cells using two cell lines, 19 and 57. Statistical significance was set at p < 0.05. C/EBPδ overexpression inhibited proliferation by accumulating cells in G1 phase of the cell cycle **F.** Anchorage independent growth assays (soft agar assays) performed on FTE-Ctrl and FTE-CEBPD-OE cells.

#### Breast and ovarian cancer cell lines

To determine C/EBPδ basal expression levels across ovarian (n = 42) and breast cancer (n = 54) cell lines, we used publically available RNA-sequence data from Medrano et al. [40]. The data uniformly showed higher C/EBPδ mRNA levels were associated with lower proliferation rates across breast and ovarian cancer cell lines (Figure 3A-B). Similarly, C/EBPδ protein expression levels in chemotherapy naive HGSC tissue by western blot analyses is low, consistent with immunohistochemical data of formalin fixed tissue (Figure 3C). To further investigate the biological consequences of C/EBPδ expression in cancer cell lines, we used three breast cancer cell lines (MCF7, T47D, MDA231), and an ovarian cancer cell line (SKOV3). MCF7 and T47D have wildtype p53 whereas MDA231 and SKOV3 have p53 mutations [41, 42]. C/EBPδ is expressed in estrogen receptor positive cancer cell lines, including MCF7 (highest), T47D and SKOV3 with no/low expression in estrogen receptor negative MDA-231 (Figure 3D). Similar to FTE cells, growth inhibitory effects of C/EBPδ overexpression were observed in cancer cell lines which resulted in reduced cell growth in MCF7, T47D, SKOV3 but an increased growth in MDA-231 compared to controls (Supplementary Figure 3A). Proliferation assays demonstrated that in MCF7, C/EBPδ overexpression decreased cell growth at day 3 (3.6-Fold, p = 0.01), day 5 (4.7-Fold, p = 0.04), day 7 (11.43-Fold, p = 0.002) and day 10 (30-Fold, p = 0.002) compared to control cells (Supplementary Figure 3B). A second breast cancer cell line, T47D, overexpressing C/EBPδ demonstrated a signifcanct decrease in cell proliferation at day 3 (11.45-Fold, p = 0.00009), day 5 (11.67-Fold, p = 0.01) and day 10 (12.5-Fold, p = 0.01) (Supplementary Figure 3B) but in the more mesenchymal cell line, MDA231, C/EBPδ increased growth rates slightly compared to controls, albeit not statistically significant (p = 0.35) (Supplementary Figure 3A, 3B). Since C/EBPδ basal expression levels were low in proliferating FTE and in HGSC, we used MCF7 cell line where basal levels of C/EBPδ can be clearly detected by western blot analysis to determine if C/EBPδ is necessary to maintain proliferation (Figure 3D). A non-targeting (NT) control cell line and a CEBPD-knockdown (KD) stable cell line were generated in MCF7 (Figure 3E). Loss of C/EBPδ expression in MCF7 had no significant effect on cell growth (Figure 3F). However, C/EBPδ over-expression resulted in a decrease of anchorage independent growth (1.6-fold, p = 0.01) (Figure 3G). In contrast, MDA231 with CEBPD overexpression resulted in increased soft agar colony formation relative to control cells (Supplementary Figure 3C). In the context of MCF7, a hormonally responsive breast cancer cell line, C/EBPδ expression is sufficient to inhibit anoikis by anchorage independent growth.

**Figure 3.**
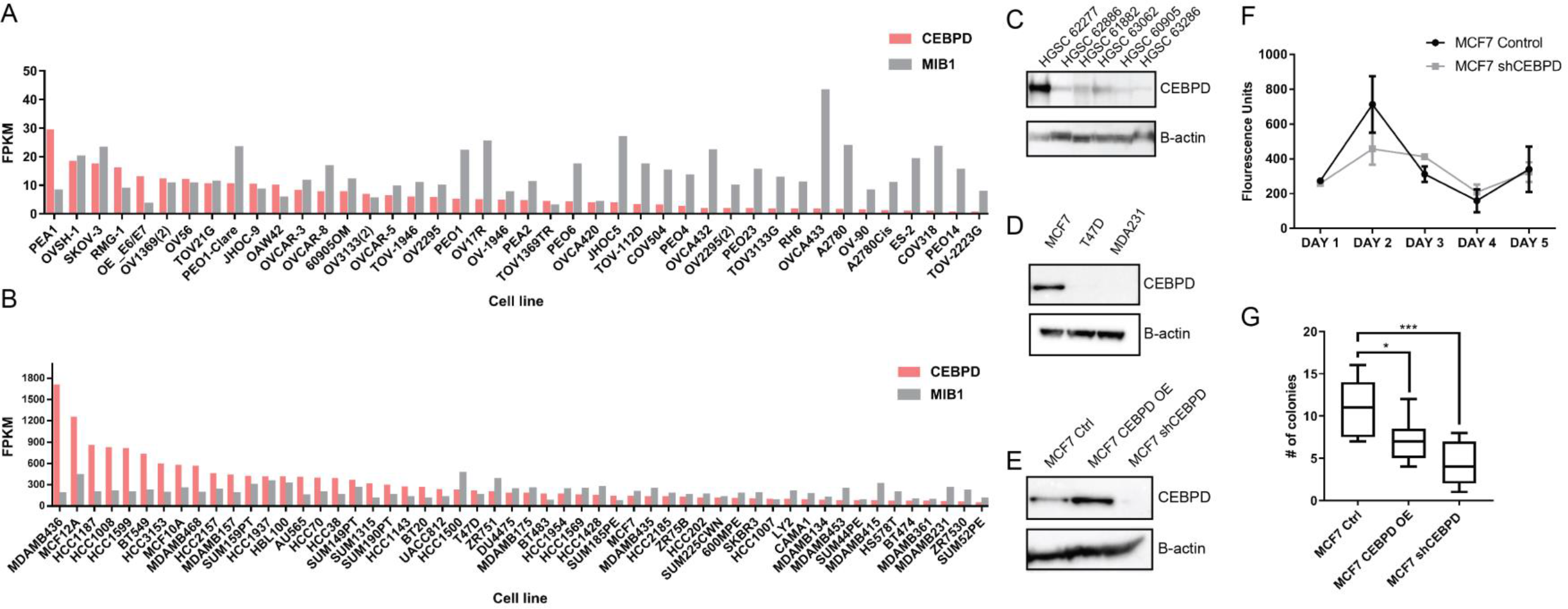
Loss of CEBPD is not necessary for proliferation in MCF7. **A.** mRNA expression levels measured as fragment per kilobase per million reads (FPKM) of C/EBPδ and MIB1 (Ki67) across a panel of 42 ovarian cancer cell lines shows an inverse correlation between C/EBPδ and MIB1 expression levels. **B.** Similarly, C/EBPδ and MIB1 mRNA expression levels were analyzed across a panel of 54 breast cancer cell lines showing higher levels of C/EBPδ were associated with lower levels of MIB1 expression. mRNA expression levels are displayed as FPKM. **C.** Western blot analysis of C/EBPδ expression across six HGSC tumour samples. B-actin is used as a loading control. **D.** Western blot analysis of C/EBPδ expression across three breast cancer cell lines, MCF7, T47D, MDA231. **E.** Western blot analysis of C/EBPδ expression in MCF7 cell line transfected with an empty GFP vector (ctrl), C/EBPδ overexpressing vector (CEBPD-OE) and a short hairpin lentiviral construct targeting C/EBPδ (shCEBPD), respectively. **F.** Cell growth was measured in MCF7-ctrl cells and MCF7-shCEBPD. **G.** Anchorage independent growth assay of MCF7 cells overexpressing C/EBPδ. P-values derived by using an unpaired t-test (*p < 0.01).

### C/EBPδ expression is associated with a MET and migratory potential in FTE and cancer

Epithelial to mesenchymal transition (EMT) has a fundamental role in cancer metastasis; restoration of the mesenchymal to epithelial transition (MET) program should efficiently slow dissemination of tumor cells [43]. Epithelial cells within fallopian tube undergo morphological changes during the menstrual cycle evident under light microscopy [44, 45]. The FTE are characteristically pseudo-stratified epithelia consisting of cuboidal and columnar epithelial cells of secretory and ciliated cells. During the pre-ovulatory (follicular) phase there is an increase in proliferation [46]. At ovulation, the secretory cells reach peak activity and secrete their nutritive contents into the lumen of the tube, then reduce in height to allow ciliated cells to move secretions by the beating of their cilia. Subsequently, in the post-ovulatory (luteal) phase, both cell types reduce in height and there is partial deciliation [47]. C/EBPδ expression in the mouse ovary was previously reported to be mediated by the luteinizing hormone, LH [48] and both mRNA and protein levels are higher in FTE in the luteal phase of the ovarian cycle [22]. Using a TMA of FTE (n = 52) [22, 46] annotated with *BRCA1* mutation and ovarian cycle status, we assessed expression of both vimentin and e-cadherin by IHC and image analysis (Figure 4A). Concurrent TMA slides were stained for all three proteins. In general, C/EBPδ basal protein expression was low. Interestingly, in the FTE luteal cases, where C/EBPδ expression was higher, we saw a downward shift in vimentin^+^ expressing FTE cells (90.2% in the follicular phase to 68.5% in the luteal phase, p = 0.33). No significant changes in e-cadherin were observed (91.0% in follicular phase to 88.9% in luteal phase, p = 0.51) (Figure 4A-B). *In vitro,* FTE with p53 mutations and hTERT retain a mesenchymal phenotype which is characterized by vimentin expression as observed in FTE *in vivo* (Figure 4D, Supplementary Figure 2B). FTE overexpressing C/EBPδ showed a distinctive epithelial-like phenotype with well-defined sheets of adhering cells compared to control cells which had more elongated mesenchymal shape (Figure 4C). Western blot analyses demonstrated a decrease in vimentin and increase in e-cadherin expression in FTE overexpressing C/EBPδ compared to controls (Figure 4D, Supplementary Figure 2B). This data suggested a role for C/EBPδ in regulating cellular phenotypes and possibly cell differentiation. Overexpression of C/EBPδ in FTE significantly increased migration compared to control cells (FTE57, 2.0-fold, p < 0.01, and FTE19, 1.6-fold, p < 0.01) (Figure 4E, Supplementary Figure 2A), but did not affect migration in one cell line (Supplementary Figure 2E). From our results, overexpression of C/EBPδ induces an MET phenotype by suppressing vimentin and increasing e-cadherin which resulted in an arrest of the cell cycle and decreased the migratory effects of cells.

**Figure 4.**
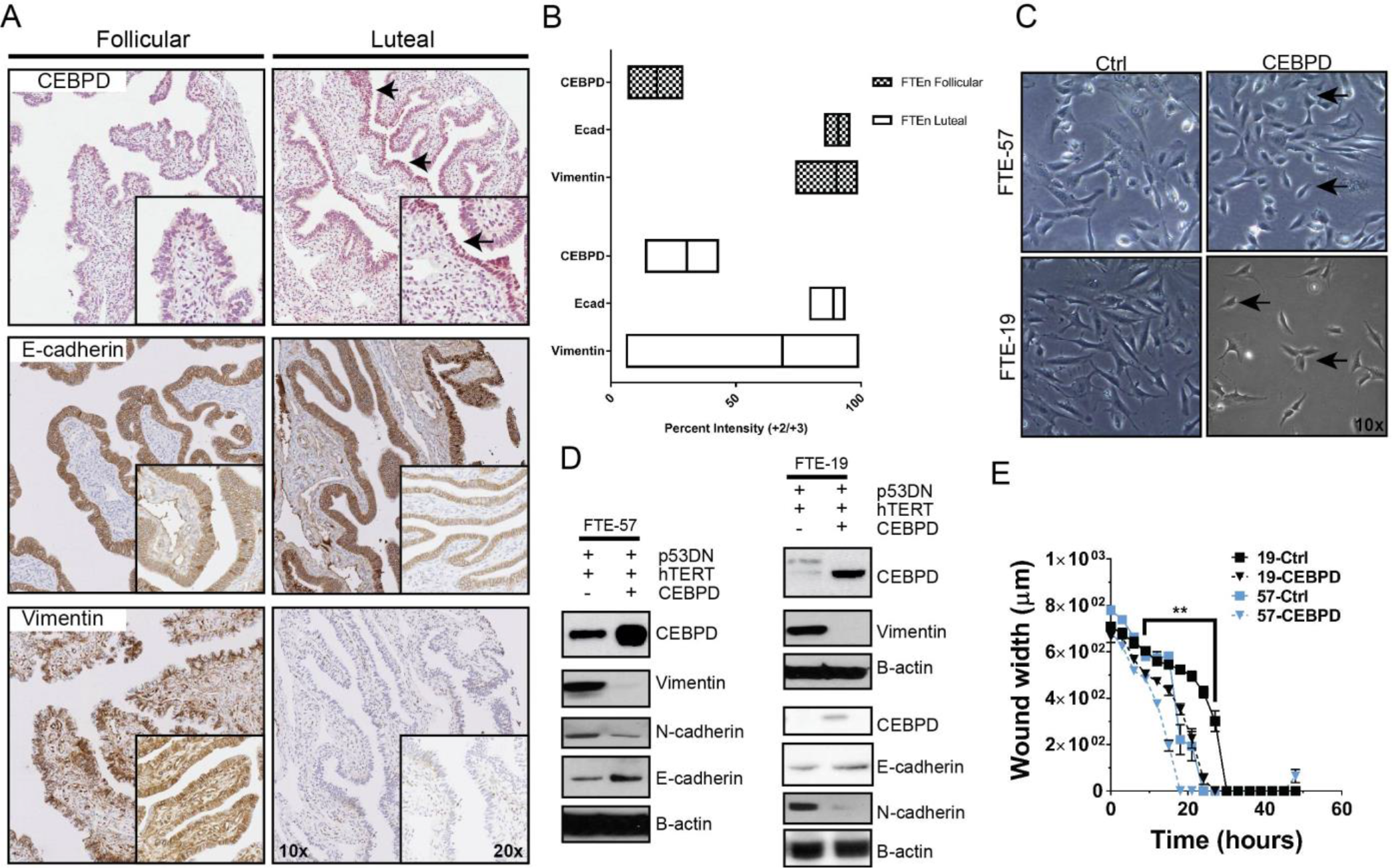
CEBPD overexpression promotes an epithelial cell phenotype with migratory potential. **A.** Representative IHC staining of C/EBPδ, E-cadherin and Vimentin in fallopian tube epithelia from the luteal and follicular phases of the menstrual cycle. **B**. Quantification of C/EBPδ, E-cadherin and Vimentin protein expression in 21 fallopian tube epithelia cases. **C**. Macroscopic images were taken using bright field light microscopy of two FTE cell models, 57 and 19 showing control verses C/EBPδ overexpressing cells. Black arrows indicate elongations from filopodia in C/EBPδ overexpressing cells. Images were taken at 20x magnification. **D.** *In-vitro* analysis of FTE-57 and FTE-19 using western blots showing C/EBPδ, e-cadherin, vimentin n-cadherin expression relative to control cells. B-actin is used as a loading control. **E.** A wound healing assay was performed on FTE-57 and FTE-19 cell lines to demonstrate migration potential. Wound width (uM) was measured against time of experiment (measured in hours).

EMT genes such as *SNAIL*, *TWIST* and the *ZEB* family of transcription factors can change the phenotypical characteristics of cells to regulate this phenomenon. E-cadherin expression has been shown to be reduced in some primary ovarian carcinomas, and then re-expressed in ovarian carcinoma effusions and at metastatic sites [49]. Further, ovarian carcinoma cells can co-express e-cadherin and the EMT-associated n-cadherin suggesting that ovarian carcinoma cells undergo incomplete EMT [50, 51]. We found similar co-expression of both e-cadherin and vimentin in benign FTE. Expression of e-cadherin goes up in neoplastic STIC lesions while the EMT associated protein vimentin is decreased (Supplementary Figure 7A-B). Furthermore, microarray analysis from a previous publication identified the heterogenous expression of multiple EMT markers in normal fallopian tube epithelia across the luteal and follicular phases (Supplementary Figure 7E). A panel of six HGSC probed for EMT/MET markers using immunoblots demonstrated the variable expression levels of these markers, consistent with other publications (Supplementary Figure 5C). IHC analysis of a STIC and HGSC sample showed that vimentin and e-cadherin levels were low in HGSC, but e-cadherin levels were higher in STIC relative to vimentin levels. (Supplementary Figure 7C, 7D). Quantitative RT-PCR (qPCR) showed a significant increase in *Snail*, *Twist* and *Zeb* in FTE-CEBPD-OE cells compared to FTE-ctrls (p < 0.01, n = 3), in the 2 independent FTE cell lines (Figure 5A). In contrast, C/EBPδ overexpression resulted in a decrease in protein expression levels of twist, slug, zeb1, vimentin and n-cadherin and an increase of snail protein compared to control cells (Figure 5B) indicating *C/EBPδ* influences a MET program in normal fallopian tube cells.

**Figure 5.**
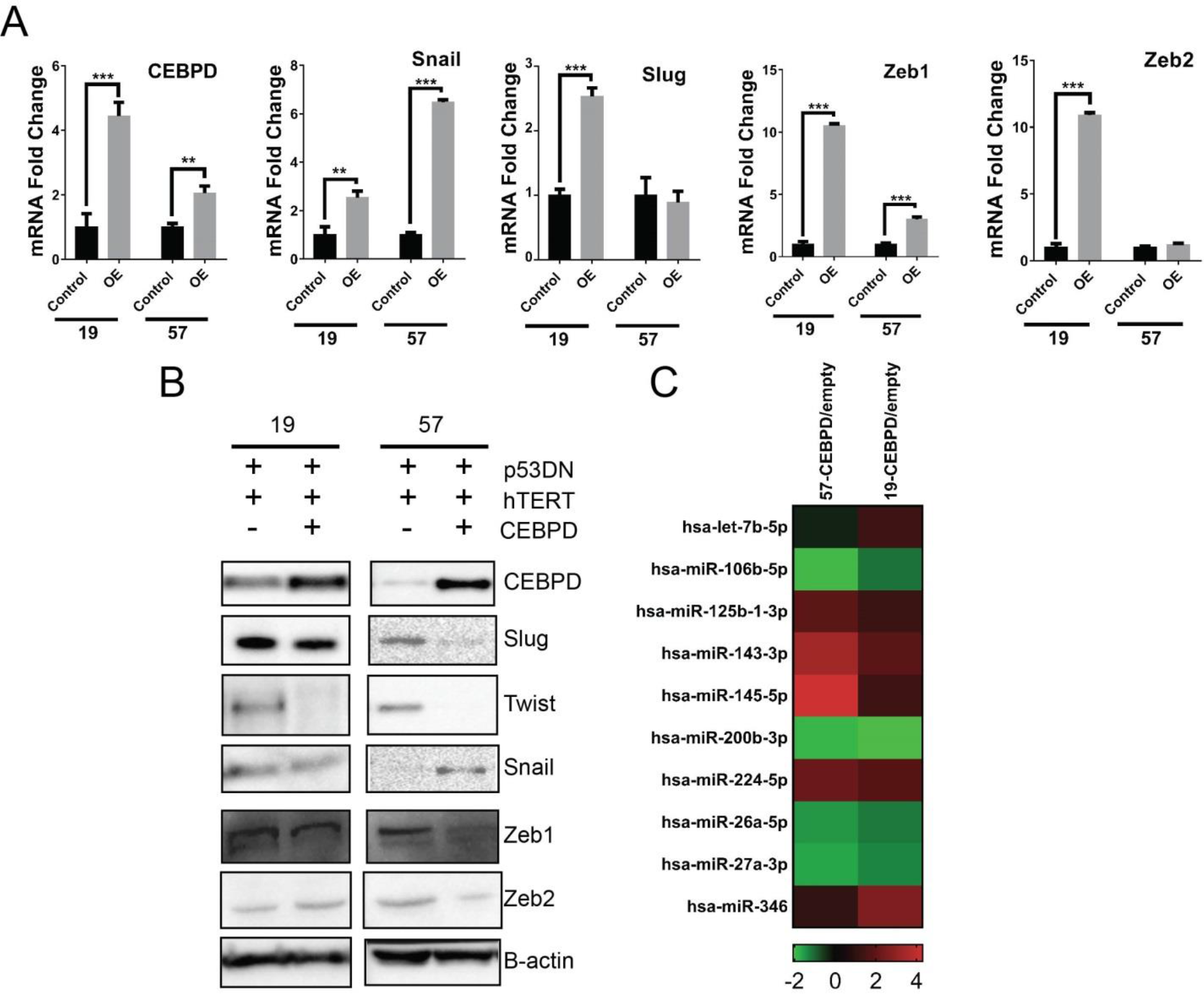
C/EBPδ overexpression downregulates EMT markers and upregulates miRNA involved with EMT downregulation. **A.** A quantitative PCR (qPCR) measured the relative mRNA expression levels of C/EBPδ, and EMT/MET markers, including Snail, Slug, Zeb1, Zeb2 in two FTE cell lines overxpresing C/EBPδ verses control cells. **B.** A western blot analysis of EMT/MET protein markers perfomed on FTE-57 and FTE-19 overexpressing C/EBPδ compared to control cells. **C.** A qPCR of miRNA was performed on both FTE-57 and FTE-19 cells showing the fold change in various miRNAs in cells overexpressing C/EBPδ compared to control cells. Upregulated miRNA (Red) and downregulated (green) were identified (p < 0.05). Data are represented as mean +/- SEM and p-values are calculated using Student t-tests, two-tailed and a 2-way ANOVA.

*In vitro*, overexpression of C/EBPδ in SKOV3 led to an epithelial phenotype characterized by well-defined cells compared to control cells, which had a more elongated, mesenchymal phenotype (Figure 6A, Supplementary Figure 4A). Furthermore, FACS analysis of SKOV3 cells demonstrated an increase in EpCAM^+^ cells and decrease in Cd49f^+^ relative to control (Supplementary Figure 4B). Across FTE cells, EPCAM^+^ cell expansion was noted, consistent with the cancer cell line (Supplementary Figure 4D). Similar to FTE, western blot analysis revealed that overexpression of C/EBPδ in SKOV3 induced e-cadherin and decreased vimentin, zeb1 and zeb2 expression levels (Figure 6B). We then studied the effect of C/EBPδ overexpression on breast cancer cells lines MCF7 and T47D which have higher overall e-cadherin compared to the more mesenchymal MDA-231 cells. C/EBPδ protein expression levels are highest in MCF7 associated with strong expression levels of epithelial markers, e-cadherin and CK8, while MDA-231 has low C/EBPδ expression which is associated with low e-cadherin and CK8 expression and high vimentin expression (Figure 6C). EMT markers such as SNAIL and TWIST were expressed in T47D cells and ZEB2 was expressed in all three cell lines (Supplementary Figure 5A). In MCF7, overexpression of C/EBPδ resulted in an increase in e-cadherin expression while silencing of C/EBPδ reversed e-cadherin levels (Figure 6D). Using T47D cells overexpressing C/EBPδ there was no effect on SNAIL and a slight decrease in TWIST expression (Supplementary Figure 5B). An in-vitro observation of MCF7 cells with overexpression C/EBPδ using bright field microscopy showed cells were more compact and rounded, indicative of an epithelial phenotype, compared to cells with a C/EBPδ knockdown, which displayed filipodia and were less compact (Figure 6E). As in FTE, the overexpression of C/EBPδ in MCF7 cells resulted in significantly more cells migrating relative to control cells (1.74 fold, p = 0.004) (Figure 6F). Taken together, as normal cells transition towards a neoplastic state, C/EBPδ expression decreases resulting in less control over the cell cycle and as a consequence pushes the cell to a more epithelial phenotype (MET) which in the context of high-grade serous cancer makes it more permissive to transformation (Figure 6G). In many *in vitro* and *in* vivo models of EMT, polarized epithelial cells EMT, cells lose polarity, cell-cell adhesion and acquire migratory and invasive properties [52]. In the FTE, a polarized epithelial layer of ciliated and secretory, cytokeratin 18^+^ cells, express both vimentin and e-cadherin and undergo a modified mesenchymal to epithelial transition during serous tubal intraepithelial transformation as seen in Supplementary Figure 6A-B.

**Figure 6.**
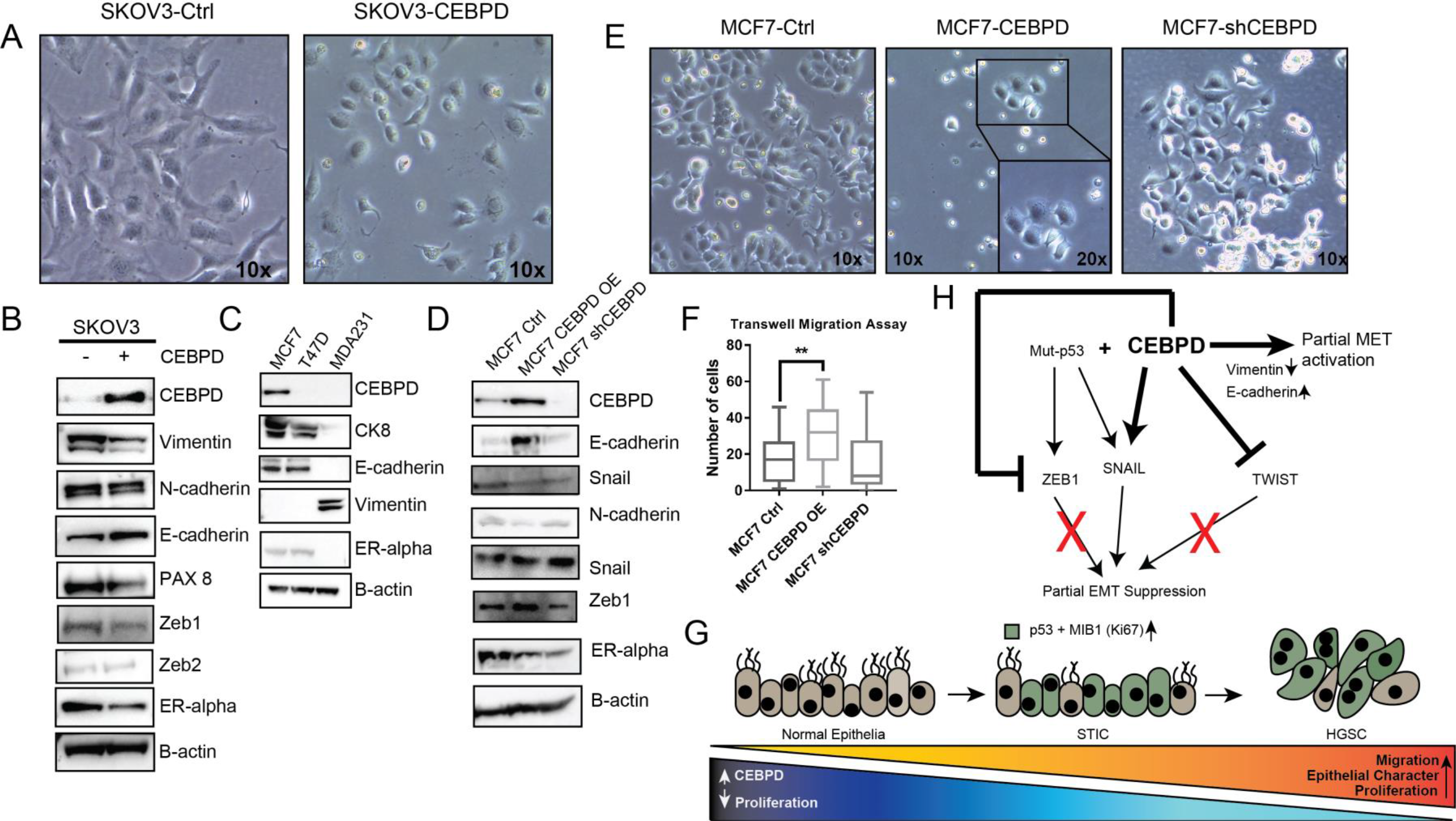
C/EBPδ increases E-cadherin expression in cancer cell lines. **A.** Representative macroscopic images of SKOV3 cell line overexpressing C/EBPδ compared to control, taken using bright field light microscopy at 20× magnification. **B**. Western blot analysis of C/EBPδ, mesenchymal markers vimentin and N-cadherin, epithelial markers E-cadherin and Pax 8, EMT/MET markers, including Zeb1, Zeb2, as well as ERα in SKOV3 cells overexpressing C/EBPδ compared to controls. **C.** CEBPD protein expression was compared in three breast cancer cell lines, MCF7, T47D, MDA 231, using western blot analysis. **D.** Western blot analysis of C/EBPδ, mesenchymal marker N-cadherin, epithelial marker E-cadherin, EMT/MET markers, including Zeb1, Snail, as well as ERα Whole-cell lysates in MCF7 control cells, C/EBPδ overexpressing cells and C/EBPδ deficient cells (shCEBPD). B-actin is used as a loading control. **E.** Images of MCF7 cells overexpressing C/EBPδ, C/EBPδ deficient cells (shCEBPD) and corresponding controls were taken using bright field light microscope at 20× magnification. **F.** Migration assay of MCF7 cells overexpressing C/EBPδ, C/EBPδ deficient cells (shCEBPD) and corresponding controls. Data are represented as mean +/- SEM and p-values are calculated using student t-tests, two-tailed. **G.** A model showing the role of C/EBPδ in HGSC development. **H.** A proposed pathway demonstrating the role of CEBPD in the EMT/MET in the context of fallopian tube epithelia.

### C/EBPδ induces an EMT regulatory phenotype

It is well established that miRNAs regulate translation efficiency of genes by targeting mRNA. [53]. This can result in differences between the amount of protein and mRNA in a cell. Certain genes, such as p53, alter the expression of miRNA which results in changes in the phenotypical characteristics of cells. Loss of p53 in mouse ovarian epithelial cells resulted in downregulation of miRNA-34b and miRNA-34c which decreased proliferation and anchorage independent growth [54]. C/EBPδ, itself has been shown to be regulated by miRNA regulatory networks with IL6 and TNF[55]. To determine whether there were differences in miRNA expression, the miRNA of FTE-CEBPD-OE was compared to FTE-ctrl cells using a miRNA qPCR Array Human Ovarian cancer array (84 miRNAs). Comparisons were made between FTE-19-CEBPD-OE/controls and FTE-57-CEBPD-OE/controls cell lines (Supplementary Figure 2A). There were 10 miRNAs that were commonly differentially expressed across both FTE-CEBPD over-expressing cells lines with a more than 2-fold change. Of these, let-7b-5p, miR-125b-1-3p, miR-143-3p, miR-145-5p, miR-224-5p and miR-346 were up-regulated and miR-26a-5p, miR-27a-3p, miR-106-5p, miR-200b-3 were down-regulated (Figure 5C, Supplementary Figure 2A). miR-200b was downregulated in cells where C/EBPδ was over-expressed. MiR-200b is downregulated in ovarian cancer and has been known to target *Zeb1*, *Zeb2* and *Slug* which are repressors of *e-cadherin*. [56, 57]. Low levels of miR-200 in addition to a decrease in e-cadherin results in an increase in vimentin, n-cadherin, twist and snail [58]. Overexpression of leb-7b suppresses cancer cell growth [59, 60] and therefore, may be one of the mechanisms by which C/EBPδ controls the fallopian tube epithelial cell cycle. C/EBPδ has also been shown to be involved in miRNA gene networks with IL6 – miR-26a-5p, TNF - miR-143-3p and miR-27a, CREBBP – miR-27b [55]. The results suggest a role for C/EBPδ in altering the miRNA expression profile within normal FTE, suggesting a plausible explanation for the discrepancy in mRNA and protein expression levels and the C/EBPδ-MET induced phenotype in the FTE cells. This data suggests a role for C/EBPδ in regulating EMT/MET in the context of p53DN fallopian tube epithelia (Figure 6H).

## Discussion

In this study, we present a role for C/EBPδ in the regulation of an EMT/MET program during early preneoplastic changes in the fallopian tube. Expression of C/EBPδ is high in ∼40% of STICs and expression from STIC to HGSC is maintained in the 40/60% high-low ratio. High expression of C/EBPδ promotes a partial EMT/MET phenotype in FTE cells. C/EBPδ has a role in promoting migration of cells despite decreasing cell proliferation, a characteristic that has also been seen in mesenchymal-like breast cancer cells expressing YB-1 [61]. It is possible that CEBPD reduces proliferation rates thereby decreasing cell density allowing for the monolayer migration [62]. In the context of a p53 mutation and CEBPD, cells become more mesenchymal relative to baseline, which results in an MET phenotype. This feature, as seen in the luteal phase of the ovarian cycle, results in slower growth. Independent of C/EBPδ, the fallopian tube epithelial cells express a mesenchymal protein, vimentin, which changes with ovulatory cycle. This highlights a new appreciation of the inherent epithelial and mesenchymal features of FTE cells which either facilitate or inhibit tumorigenesis (Figure 7G,7H).

Previously, by comparing hormonally driven changes of phenotypically normal FTE from women with and without a BRCA1 mutation, we identified several differentially expressed genes with known functions that promote tumor development and metastasis [22, 63]. Amongst these genes, C/EBPδ, a transcriptional regulator of cellular differentiation, inflammatory signaling, hypoxia adaptation and metastatic progression, was increased during the luteal phase of the ovarian cycle [22]. C/EBPδ has dualistic roles as a tumor suppressor or oncogene and has been called a master regulator gene. In breast cancer cell line, MCF7, C/EBPδ was shown to decrease cylinD1 by mediating CCND1 degradation through Cdc27/APC3 regulation [64, 65]. In the epidermoid skin cancer cell line, A431, C/EBPδ also decreased cancer cell proliferation induced by interacting with E2F1 and Rb [31]. In leukemia, CML, KCL22 and K562 cell lines, expression of C/EBPδ was associated with downregulation of c-Myc and cyclin E and upregulation of the cyclin-dependent kinase inhibitor p27 [28]. We sought to establish C/EBPδ’s role in the pathogenesis of high-grade serous ovarian cancer. Our tissue-based data revealed that 77% of a cohort of HGSC cases had low C/EBPδ protein expression compared to LGSCs. This difference may be reflective of the underlying pathogenic routes to these distinct tumors that arise in the fallopian tube [2, 66]. HGSC is fast growing and has a high proliferative index compared to LGSC, which is slow growing and indolent [67]. In our cohort of normal FTE and serous intraepithelial carcinoma *in situ* (STIC) cases, C/EBPδ expression was inversely correlated to Ki67 expression. Higher expression levels of C/EBPδ were associated with lower Ki67 staining and the trend was consistent in HGSCs. This suggests that C/EBPδ is preferentially downregulated in highly proliferative tissues of the fallopian tube and cancer.

Since C/EBPδ expression is higher in the normal tissues compared to 70% of HGSC, we used an *in vitro* FTE model to explore the effects of C/EBPδ overexpression in p53 dominant negative mutant cells [6]. In this study, C/EBPδ overexpression decreased cellular growth resulting in accumulation of the cells in the G1/S phase of the cell cycle. This observation was extended to ovarian and breast cancer cell lines, suggesting C/EBPδ is enough to inhibit growth. Unfortunately, endogenous levels of C/EBPδ were too low to observe an additional decrease in endogenous protein levels in the modeled premalignant fallopian tube cells. However, silencing C/EBPδ in MCF7 did not affect growth, suggesting that C/EBPδ is sufficient but not necessary for proliferation.

Within the normal tissue, C/EBPδ levels are higher in the luteal phase of the menstrual cycle relative to the follicular phase. The luteal phase constitutes a highly inflammatory milieu with fallopian tube epithelial cells undergoing differentiation to accommodate the motility of the ovum toward the ampulla for impregnation. C/EBPδ is rapidly induced by inflammatory signals, cytotoxic factors and stressful conditions, which are characteristics of the luteal phase [23, 68]. Given that C/EBPδ has a dichotomous role in regulating proliferation and differentiation [28], we hypothesized the expression of C/EBPδ might be slowing the growth of cells to prepare them for a transitional state. Indeed, we describe C/EBPδ’s role for the first time in mediating an EMT/MET switch thereby providing FTE with epithelial plasticity and enabling the migratory potential of these cells. We hypothesize higher levels of C/EBPδ in premalignant FTE might alter cell phenotype toward an epithelial fate with potential to migrate to the ovarian surface epithelium.

One of the main points of debate regarding the fallopian tube epithelial origin of HGSC is how mesenchymal cells of origin (i.e. fallopian tube epithelial cells) form serous epithelial ovarian carcinoma. Factors other than location and proximity contribute to seeding to the ovary by fallopian tube precursor lesions. We hypothesized menstrual cycle associated genes with dual role of controlling proliferation and differentiation might play an important role in this process. In our FTE cell culture model, a p53 dominant negative (R175H) mutation was introduced to recapitulate one of the earliest known events in HGSC, namely, the p53 signature. Loss of p53 was previously shown to induce mesenchymal-like features in normal mammary epithelial cells [56]. We report similar findings in FTE. We also observed expression of twist, slug, zeb1 and zeb2 at the protein level in p53 deficient FTE cells, a feature consistent with a mesenchymal phenotype [69].

Here we showed C/EBPδ overexpression induced a partial MET, characterized by an increase in expression of epithelial markers, E-cadherin and CK7, and a decrease in expression of mesenchymal markers, including vimentin and n-cadherin. This was observed in FTE precursor lesions as well as cancer cell lines. Furthermore, upon C/EBPδ overexpression, a decrease in the expression of twist, slug and zeb1 were observed at the protein level. The data suggests a role for C/EBPδ in regressing the mesenchymal phenotype of p53 mutated FTE cells. The overexpression of C/EBPδ still increased the migratory rate of FTE cells and cancer cell lines. However, knocking down C/EBPδ in a cancer cell line did not demonstrate a significant difference in migration relative to controls. In urothelial cancers, C/EBPδ enhanced the invasiveness of UC cells through direct binding and upregulation of MMP*2* [70]. This highlights a dichotomous role of C/EBPδ as a tumor suppressor or oncogene with tumor promoting potentials depending on the tissue and the genomic context within which the gene is expressed. Primary fallopian tube cells have limited growth expansions in vitro [6] and therefore, conducting these experiments in the p53 wild type setting was not feasible. However, the presence of p53-R175H mutation along with C/EBPδ over-expression promotes cellular migration.

Human Fallopian tube epithelial cells do not require intravasation to metastasize to the ovary and other organs in the abdominal cavity. FTE cells are exposed to the peritoneal cavity where they can slough off easily and disseminate to other locations. This study highlights new roles for C/EBPδ in HGSC and separately features intriguing EMT/MET hybrid FTE phenotypes influenced by the ovulatory cycle. This hybrid-pleiotropic phenotype in cancer is implicated in cancer metastasis, cancer stem cell plasticity, chemoresistance and immune evasion [71, 72], all features also attributed to ovarian cancer biology and heterogeneity. These EMT/MET hybrid protein expressions observed in the histologically normal fallopian tube epithelia and pre-malignant lesions need further study to identify core transcriptional and genomic/epigenomic networks driving fallopian tube epithelia transformation. Further, FTE cells endogenously express well established mesenchymal markers like vimentin and epithelial markers as e-cadherin simultaneously. Acquisition of mutant p53 is not required for these cells to express mesenchymal genes but may play a role in promoting anoikis and survival in the peritoneal cavity once detached from the basement membrane. During the ovulatory cycle, C/EBPδ decreases cell growth and can influence mesenchymal gene expression potentially through miRNAs. Although the connection between C/EBPδ and miRNA requires further study, several publications have highlighted the role of C/EBPδ in regulating miRNA resulting in downstream phenotypical differences in cells with and without C/EBPδ, as is the case of miR-193b in urothelial human carcinoma’s which demonstrated and anti-tumorigenic function of C/EBPδ through miR-193b in NUTB1 human urothelial carcinoma cell line [73]. In this context, we suggest a model whereby C/EBPδ’s initial expression in premalignant FTE would slow cell cycle progression and allow these cells to acquire epithelial phenotype with potential to avoid anoikis and migrate to the ovary. Once these cells have migrated to the ovary, decreased expression of C/EBPδ promotes the proliferative potential of the cells which maintain a mesenchymal phenotype reminiscent of the cell of origin. Future studies are warranted to investigate which other genes and/or gene networks promote these MET/EMT switches in early fallopian tube epithelial cells transformation and whether these networks are elicited in chemoresistant ovarian cancer.

## Material and Methods

### Case collection

The University Health Network Research Ethics Board approved the study protocol for collection of tissue and clinical information for all patients. Each patient provided written informed consent allowing for the collection and use of tissue for research purposes. H&E sections of high-grade serous carcinoma, borderline and low-grade serous carcinoma were reviewed by a gynecological pathologist (P.S.) prior to use in the study. Diagnosis of each case was retrieved from the UHN ovarian tissue bank prior to review. To validate immunohistochemical protein expression of samples, whole sections of tissue were cut from formalin fixed paraffin embedded tissue and analyzed as in our previous publication [6, 13, 74]. A previously published cohort (n = 15) of serous tubal intraepithelial carcinoma (STIC) cases using Abcam morphological and immunohistological features including cellular crowding, loss of nuclear polarity and presence of p53 and Ki67, from women having adnexal high-grade serous carcinoma [6, 13, 46].

### Immunohistochemistry

Immunohistochemistry was performed using standard procedures as previously described [63] with the following modifications. The following antibodies were used at these dilutions: Ki67 (Lab Vision, Thermo Scientific, Waltham, Massachusetts, USA) 1/1000; p53 (Novocastra, Leica, Wetzlar, Germany) 1/200; C/EBPδ 1/200 (sc-636) (Santa Cruz Biotechnology, Inc., Dallas, Texas, USA); E-cadherin 1/100 (AB15148) (Abcam, Cambridge, United Kingdom); Vimentin 1/100 (5741S) (Cell Signaling Technology, Danvers, Massachusetts, USA). Appropriate negative and positive controls were performed to determine specificity of antibodies. Stained slides were scanned using the ScanScope XT slide scanner (Aperio Technologies, Inc, Leica) to create digital images at 40X magnification which were then quantified for intensity and percentage of cells staining using a nuclear algorithm as previously described (Spectrum Plus, Image Analysis Toolbox, TMALab II, Aperio, Inc.) [22, 74]. Intensity levels were based on an absorption range scale of 0-255 (0 = black; 255 = white). Weak Intensity Staining (1+) ranged from 200-215; medium intensity staining (2+) from 180-200; and strong intensity staining (3+) from 0-180. Intensity levels + 2 and + 3 were combined to create a composite score for the percent positive nuclei present in each case. Images were annotated to include only epithelium while excluding stroma.

### Immunofluorescence

Cells were grown in 6-well plates (Falcon) coated with collagen IV and fixed with 4% PFA for 5 minutes, permeabilized with 0.3% Triton-X/PBS then blocked with 5% goat serum (Gibco, Life technologies) in PBS. Primary antibodies: C/EBPδ (SC-636), CK18 (M701029) (Dako, Agilent Technologies, Santa Clara, California, United States), Pax8 (10336-1-AP) (ProteinTech, Rosemont, IL, USA), E-cadherin (AB15148) (Abcam), Vimentin (5741S) (Cell Signaling), were applied overnight at 4 °C. Primary antibody was removed by washing samples with 1x PBS three times and was incubated with appropriate fluorophore-labeled secondary antibodies (Jackson ImmunoResearch Laboratories, West Grove, PA, USA) and Vectashield (H-1200, Vector Laboratories, Burlingame, CA, USA) in a dark area for 45 min. Cells were again washed for three times with 1x PBS and was mounted onto a coverslip and dried in the dark for 10 min. Samples were visualized using a Leica Axioimager (Leica).

### Fallopian tube epithelia tissue cultures

Surgical samples were obtained from the University Health Network with patient consent and Research Board Ethics Approval. In brief, fimbriae were collected after prophylactic hysterectomy or salpingo-oophorectomy and incubated for 4-16 hours at 37°C in pronase and subsequently cultured as previously described [6]. Cells were immortalized via infection with a lentiviral dominant negative TP53 (R175H) vector and retroviruses human telomerase (hTERT) [6]. Cells were infected with a lenti-viral C/EBPδ overexpression construct 24 to 48 hours after cells reached 70% confluence. Each plate was grown till confluence and harvested for different molecular assays. For biological replicates, all molecular analyses of cell lines for RNA and protein extraction were performed from the cell population. That is, at time of collection, cells were divided into two pellets. Additionally, for FACS and immunofluorescence, the same population of cells were pelleted for RNA and protein assays.

### Cell lines

SKOV3, OVCAR3, MCF7, MDA231, T47D were obtained from the ATCC (Manassa, Virginia, USA). SKOV3 (ATCC-HBT-77) was grown in McCOY’s 5a (Life technologies, Carlsbad, California, USA) supplemented with 10% fetal bovine serum (FBS, Wisent technologies, St-Bruno, Quebec Canada). MCF7 (ATCC-HTB-22) and MDA231 (ATCC-HTB-26) cells were grown in DMEM/F12 (Life technologies), supplemented with 10% FBS. T47D (ATCC-HTB-133) and OVCAR3 (HTB-161) cells were grown in RPMI 1640 (Life technologies), supplemented with 10% and 20% FBS, respectively. Culture method was followed as described by ATCC for each cell line.

### Virus Production and Infection

HEK-293 T cells were seeded at a density of 1.0 x10^∧^6 cells per plate in a 6 cm tissue culture dish overnight with low antibiotic growth media (DMEM + 10% FBS). Cells were incubated until 70% confluence was achieved. A mixture of 3 transfection plasmids were produced by combining 2 µg pMDG.2 (Addgene #12259) (Addgene, Watertown, Massachusetts, USA); 4 µg of pCMV delta R8.2 (Addgene #12263) and 5 µg of vector (pCDH-CMV-MCS-EF1-GFP empty, pCDH-CMV-MCS-EF1-GFP-CEBPD - CEBPD-OE). As per manufacturer’s protocol, GenJet DNA In-vitro Transfection reagent (Ver. II) (SignaGen Laboratories, Rockville, MD, USA) was used as the transfection reagent. Reagents were added to each plate drop-wise and was incubated overnight (16 hours, 37°C, 5% CO_2_). Next day, media was subsequently removed and replaced with high BSA growth media and again incubated for 24 hrs. Supernatant was then collected and passed through a 0.45um filter and snap frozen and stored at −80°C until required for use. A second round of collection was performed 72 hours after virus transfection and was also filtered and stored at −80°C.

### Protein isolation, western blot and antibodies

Cell samples were washed with 1x PBS and trypsinized using 0.25% trypsin dissociation reagent (Life technologies) for 5 min at 37 degree Celsius. TNS was added to samples and was spun at 1000 rpm for 5 minutes. Supernatant was aspirated, and samples were placed on ice before lysing. Cells were lysed with NP40 buffer (Life technologies) and protease inhibitor cocktail (Thermo Scientific). Samples were incubated for 15 min on ice and subsequently clarified by centrifugation (14000rpm for 30min). The supernatant was collected and quantified using BioRad DC Protein Assay (BioRad Laboratories, Hercules, California, United States). Samples were re-suspended in 4x LDS sample buffer (Life technologies), boiled and 30ug of protein was resolved by SDS-Page 4-12% Bis-tris Gels (Novex, Life technologies). Protein samples were transferred to BioRad PVDF membrane and were blocked with 5% milk powder dissolved in 1x TBS-T for one hour or overnight. Primary antibodies were diluted in blocking buffer (5% skimmed milk, Nestle, Vevey, Switzerland) and incubated with membranes overnight at 4°C. Membranes were washed for 10min, three times and probed with secondary antibody, washed three times for 15min using 1x TBS-T and developed using ECL Prime western blotting detection reagent (GE Healthcare Life Sciences, Pittsburgh, PA, USA). Each membrane was imaged using BioRad ChemiDoc and images analyzed using BioRad Image Lab software (BioRad). The following antibodies were used in western blots: C/EBPδ (Santa Cruz, SC-636, 1/200), B-actin (Sigma, A2228, 1:1000) (Sigma-Aldrich, St. Louis, Missouri, USA), P53 (Santa Cruz; sc-126, 1/500), PAX8 (ProteinTech, 10336-1-AP, 1/1000), Snail1 (Cell Signaling, 3895S, 1/500), Slug (Cell Signaling, 9585S, 1/500) Vimentin (Cell Signaling, 5741S, 1/500), E-cadherin (Abcam, AB15148 1/500), N-cadherin (SantaCruz, sc-7939, 1/500), SIP1 (Santa Cruz, sc-48789, 1/200), Twist (Santa cruz, sc-81417, 1/500) and goat HRP-IgG (anti-mouse or anti-rabbit) (Santa Cruz, 1:10,000).

### Cloning Strategy

Overexpression vectors were constructed using pCDH-CMV-MCS-EF1-GFP cloning/ expression vector (System Biosciences, Palo Alto, CA,USA). C/EBPδ insert was cut out of a PCR4 TOPO cloning vector (Invitrogen, Carlsbad, California, USA) using EcoRI (New England Biolabs (NEB), Ipswich, Massachusetts, USA). The cDNA was then ligated into pCDH-CMV-MCS-EF1-GFP using Takara ligation Kit (Clontech, Mountain View, California, USA) and was confirmed in the expression vector by DNA sequencing at The Centre for Applied Genomics (SickKids, Toronto, Ontario, Canada). Lentivirus was produced using HEK-293T cells (Clontech) and the overexpression vector was confirmed in the cells by GFP fluorescence and western blot (Figure 2A-B, Supplementary Figure 4A).

### Fluorescence Activated Cell Sorting (FACS)

For cell cycle regulation assessment, cells were washed in 1x PBS and 10uM BrdU-APC was added to cells in a dark environment. Plates with BrdU were then incubated at 37 degrees for 4 hrs. Cells were then washed twice with ice cold PBS, trypsinized with 0.25% trypsin dissociation reagent and neutralized for counting. Nuclei preparation and staining was performed by adding 0.08% Pepsin and 2N HCl and IFA/0.5% Tween20 (Sigma). Samples were incubated in the dark ad 100ul anti-BrdU-APC was added while samples were incubated on ice. 5ug/ml of propidium iodide was added to cells and incubated on ice for 15min. Flow cytometry was carried out on BD FACS Calibur (BD biosciences, San Jose, CA, USA). Data was analyzed using FlowJo v10 (Flowjo LLC, Ashland, OR, USA). Cell surface staining was performed on single cell suspension. EpCAM-PE (Life technologies, VU-1D9) and CD49f-APC (FAB13501A) (R&D Biosystems, Minneapolis, Minnesota, USA) was added to the cell suspension and incubated in the dark for 40-60min. Cells were then washed and centrifuged at 400g for 5 min at room temperature and subsequently vortexed to dissociate pellet. Stained cells were re-suspended in staining buffer and flow cytometry acquired on BD Calibur flow cytometer (BD biosciences).

### Soft Agar Assay

Base agar (1% agarose Difco Agar Noble, BD biosciences) was added to 6 well plates and allowed to solidify for 5 min. Top agarose layer was made by combining 0.7% agarose with media. 5000 cells were added to the mixture and plated on 6 well plate. Cells were kept in an incubator at 37°C for 14 days. Cells were fed twice a week. After 14 days, each plate well was stained with 0.5ml of crystal violet (0.005%) for approximately 1 hr. Plates were then imaged using a dissecting microscope and camera setup (Leica). Images were imported into Image J (National Institute of Health, Bethesda, Maryland, USA) and analyzed.

### Quantitative PCR

RNA was isolated from FTE cells lysed with Trizol reagent (Invitrogen). 1ug of total RNA was reverse-transcribed using qScript cDNA SuperMix (Quanta Biosciences, Beverly, MA, USA). Real-time quantitative PCR (RT-qPCR) was performed using PerfeCTa Sybr Green FastMix, Rox (Quanta Biosciences) according to the manufacturer’s protocol using the ABI PRISM 7900HT Sequence Detection System (Applied Biosystems, Foster City, California, USA). The target CT values were normalized to b-actin. The N-fold differential expression was assessed using the 2-^(ΔΔCT)^ method to determine differences between treated cells and controls. Primer sequences were obtained from PrimerBank [75, 76]. qRT-PCR as mean ± s.e.m. Student’s t-test with n = 3, p ≤ 0.05 unless noted otherwise in text. N = 3 represents 3 independent experiments with 3 technical replicates within each experiment.

### miRNA Assay

FTE cells were harvested from 6 cm plates and RNA was isolated using miRNAeasy micro kit (Qiagen, Hilden, Germany). Quantity and quality of total RNA was analyzed using Nanodrop (Thermo Scientific). RNA was reverse transcribed into cDNA using miScript II RT kit (Qiagen). cDNA was then processed using miScript SYBR Green PCR kit (Qiagen) and was run on miScript miRNA PCR Array Human Ovarian Cancer plates (Qiagen, MIHS-110ZE-4, 384 well plate). PCR plates were read the ABI PRISM 7900HT Sequence Detection System (Applied Biosystems). Results were outputted with 2-ΔCt values for each gene in each treatment group compared to the control group (n = 3). Qiagen software was used to analyze results and student’s t-test provided statistically significant miRNA (p < 0.05).

### Wound-healing Assay

63857 p53DN-hTERT, 3619 p53DN-hTERT cells were seeded in 96-well plates (Essen ImageLock) and grown to confluence. Scratch wounds were generated using the 96-pin WoundMaker (Essen BioScience, Ann Arbor, MI, USA) and wells were gently washed with 1xPBS to remove non-adherent cells. Cells were imaged using the INCUCYTE™ Kinetic Imaging System (Essen BioScience) and wound width was determined using the INCUCYTE™ cell migration software module.

### Migration Assay

MCF7 cells and FTE cells were seeded onto Costar Transwell migration assay (Corning Costar 3472) (Corning Inc., Corning, New York, USA) at a density of 50000 cells. Plated cells were initially grown in serum free media for 24 hrs. After trypsinization using Trypsin (EDTA 00.25%) (Life Technologies), cells were counted and plated onto the Transwell migration assay which was then placed in DMEM/F12 + 10%FBS for 12 hours. Transwell’s were then removed from assay and washed PBS and H2O to remove unbound cells. 1% Crystal Violet + 2% ethanol was added to the Transwell and allowed to incubate at room temperature for 15 minutes at which point the wells were rinsed with H2O and dried. Each well was imaged using an inverted microscope (Leica) and cells were counted in ImageJ (Image J labs). Results were analyzed and graphed using GraphPad graphing software (GraphPad, La Jolla California USA)

### Proliferation Assay

For Proliferation Assay, 3.0 × 10^4^ cells were seeded onto a 6-well plate (Falcon, Corning Inc.) and plated with 2 ml of fresh media. Cells were counted every 1-3 days using CyQuant Direct Proliferation Assay kit (Life technologies). An inverted plate reader (Flexstation 3) (Molecular Devices, San Jose, California, USA) was used to quantify the amount of fluorescence emitted by live cells and provide a count of live cells present. Softmax Pro software (Molecular Devices) was used to analyze the results.

### Statistical Analysis

Statistical analysis was performed using GraphPad Prism Software (GraphPad). The Log-rank test was used in Kaplan-Meir, Mantel-Cox regression analysis to compare C/EBPδ expression on overall survival and progression free survival. MicroRNA analysis was performed using Qiagen SaBiosciences miScript miRNA PCR Array Data Analysis Software (Qiagen). Values are expressed as means ±SEM and were performed three times, t-tests were performed to determine statistical significance, unless otherwise noted. A value with p < 0.05 was considered to be statistically significant.

## Acknowledgments

We thank the UHN Biobank and SCCC Biospecimen Shared Resource for sample acquisition, UHN Cancer Biobank Core Laboratory and University of Miami Confocal Core for digital imaging and image analysis services. This study was funded by the CDMRP Ovarian Cancer program (W81WH-0701-0371, W81XWH-18-1-0072), the Princess Margaret Cancer Centre Foundation, Foundation for Women’s Cancer – The Belinda-Sue/Mary-Jane Walker Fund, Colleen’s Dream Foundation and Sylvester Comprehensive Cancer Center.

## Author contribution

Conception and design: SG, RS and PS. Development of methodology: RS, RC and SG; Acquisition of data: SG, RS, MB, LD, AM; Analysis and interpretation of data: RS, RC, BS, PS and SG; Writing, review and editing: all authors.

## Reference

1. Bowtell, D.D., et al., Rethinking ovarian cancer II: reducing mortality from high-grade serous ovarian cancer. Nat Rev Cancer, 2015. 15(11): p. 668–79.

2. Vang, R., M. Shih Ie, and R.J. Kurman, Fallopian tube precursors of ovarian low- and high-grade serous neoplasms. Histopathology, 2013. 62(1): p. 44–58.

3. Piek, J.M., et al., Women harboring BRCA1/2 germline mutations are at risk for breast and female adnexal carcinoma. Int J Gynecol Pathol, 2003. 22(3): p. 315–6; author reply 315-6.

4. Piek, J.M.J., et al., Dysplastic changes in prophylactically removed Fallopian tubes of women predisposed to developing ovarian cancer. The Journal of Pathology, 2001. 195(4): p. 451–456.

5. Folkins, A.K., et al., A candidate precursor to pelvic serous cancer (p53 signature) and its prevalence in ovaries and fallopian tubes from women with BRCA mutations. Gynecol Oncol, 2008. 109(2): p. 168–73.

6. George, S.H., et al., Loss of LKB1 and p53 synergizes to alter fallopian tube epithelial phenotype and high-grade serous tumorigenesis. Oncogene, 2015.

7. George, S.H. and P. Shaw, BRCA and Early Events in the Development of Serous Ovarian Cancer. Front Oncol, 2014. 4: p. 5.

8. Karst, A.M., K. Levanon, and R. Drapkin, Modeling high-grade serous ovarian carcinogenesis from the fallopian tube. Proceedings of the National Academy of Sciences of the United States of America, 2011. 108(18): p. 7547–52.

9. Jazaeri, A.A., et al., Molecular requirements for transformation of fallopian tube epithelial cells into serous carcinoma. Neoplasia, 2011. 13(10): p. 899–911.

10. Sehdev, A.S., et al., Serous tubal intraepithelial carcinoma upregulates markers associated with high-grade serous carcinomas including Rsf-1 (HBXAP), cyclin E and fatty acid synthase. Mod Pathol, 2010. 23(6): p. 844–855.

11. Visvanathan, K., et al., Diagnosis of serous tubal intraepithelial carcinoma based on morphologic and immunohistochemical features: a reproducibility study. The American journal of surgical pathology, 2011. 35(12): p. 1766–75.

12. Shaw, P.A., et al., Candidate serous cancer precursors in fallopian tube epithelium of BRCA1/2 mutation carriers. Modern Pathology, 2009. 22(9): p. 1133–1138.

13. Milea, A., et al., Retinoblastoma pathway deregulatory mechanisms determine clinical outcome in high-grade serous ovarian carcinoma. Mod Pathol, 2013.

14. Levanon, K., et al., FOXO3a loss is a frequent early event in high-grade pelvic serous carcinogenesis. Oncogene, 2014. 33(35): p. 4424–32.

15. Norquist, B.M., et al., The molecular pathogenesis of hereditary ovarian carcinoma: alterations in the tubal epithelium of women with BRCA1 and BRCA2 mutations. Cancer, 2010. 116(22): p. 5261–71.

16. Cancer Genome Atlas, N., Integrated genomic analyses of ovarian carcinoma. Nature, 2011. 474(7353): p. 609–15.

17. Karst, A.M., et al., Cyclin e1 deregulation occurs early in secretory cell transformation to promote formation of fallopian tube-derived high-grade serous ovarian cancers. Cancer Res, 2014. 74(4): p. 1141–52.

18. Karst, A.M., et al., Stathmin 1, a marker of PI3 K pathway activation and regulator of microtubule dynamics, is expressed in early pelvic serous carcinomas. Gynecologic oncology, 2011. 123(1): p. 5–12.

19. Kuhn, E., et al., Telomere length in different histologic types of ovarian carcinoma with emphasis on clear cell carcinoma. Mod Pathol, 2011. 24(8): p. 1139–45.

20. Lengyel, E., Ovarian cancer development and metastasis. Am J Pathol, 2010. 177(3): p. 1053– 64.

21. Sodek, K.L., et al., Cell-cell and cell-matrix dynamics in intraperitoneal cancer metastasis. Cancer Metastasis Rev, 2012. 31(1-2): p. 397–414.

22. George, S.H., et al., Identification of abrogated pathways in fallopian tube epithelium from BRCA1 mutation carriers. The Journal of pathology, 2011. 225(1): p. 106–17.

23. Ramji DP, F.P., CCAAT/enhancer-binding proteins: structure, function and regulation. Biochem J., 2002. 365(Pt 3): p. 561–75.

24. Sterneck, K.B.a.E., The Many Faces of C/EBPδ and their Relevance for Inflammation and Cancer. Int J Biol Sci., 2013. 9(9): p. 917–933.

25. Wu, S.-R., et al., CCAAT/Enhancer-binding Protein δ Mediates Tumor Necrosis Factor α-induced Aurora Kinase C Transcription and Promotes Genomic Instability. The Journal of Biological Chemistry, 2011. 286(33): p. 28662–28670.

26. Carro, M.S., et al., The transcriptional network for mesenchymal transformation of brain tumours. Nature, 2010. 463(7279): p. 318–25.

27. Cooper, L.A.D., et al., The Tumor Microenvironment Strongly Impacts Master Transcriptional Regulators and Gene Expression Class of Glioblastoma. The American Journal of Pathology, 2012. 180(5): p. 2108–2119.

28. Gery, S., et al., C/EBPδ expression in a BCR-ABL-positive cell line induces growth arrest and myeloid differentiation. Oncogene, 2005. 24: p. 1589.

29. Sanford, D.C. and J.W. DeWille, C/EBPdelta is a downstream mediator of IL-6 induced growth inhibition of prostate cancer cells. Prostate, 2005. 63.

30. Hutt, J.A. and J.W. DeWille, Oncostatin M induces growth arrest of mammary epithelium via a CCAAT/enhancer-binding protein delta-dependent pathway. Mol Cancer Ther, 2002. 1.

31. Pan, Y.-C., et al., CEBPD reverses RB/E2F1-mediated gene repression and participates in HMDB-induced apoptosis of cancer cells. Clinical Cancer Research, 2010.

32. Ko, C.-Y., et al., Epigenetic Silencing of CCAAT/Enhancer-binding Protein δ Activity by YY1/Polycomb Group/DNA Methyltransferase Complex. Journal of Biological Chemistry, 2008. 283(45): p. 30919–30932.

33. Tang, D., G.S. Sivko, and J.W. DeWille, Promoter methylation reduces C/EBPδ (CEBPD) gene expression in the SUM-52PE human breast cancer cell line and in primary breast tumors. Breast Cancer Research and Treatment, 2006. 95(2): p. 161–170.

34. Takayuki Ikezoe, S.G., Dong Yin, James O’Kelly, Lise Binderup, Nathan Lemp, Hirokuni Taguchi and H. Phillip Koeffler, CCAAT/Enhancer-Binding Protein δ: A Molecular Target of 1,25-Dihydroxyvitamin D3 in Androgen-Responsive Prostate Cancer LNCaP Cells. Cancer Res, 2005. 65(11): p. 4762–4768.

35. Kuppusamy Balamurugan, S.S., Kimberly D. Klarmann, Youhong Zhang, Vincenzo Coppola, Glenn H. Summers, Thierry Roger, Deborah K. Morrison, Jonathan R. Keller & Esta Sterneck, FBXW7α attenuates inflammatory signalling by downregulating C/EBPδ and its target gene Tlr4. Nature Communications, 2013. 4(1662): p. 1–12.

36. Gigliotti, A.P., et al., Nulliparous CCAAT/enhancer binding protein delta (C/EBPdelta) knockout mice exhibit mammary gland ductal hyperlasia. Exp Biol Med, 2003. 228.

37. Thangaraju M, R.M., Bierie B, Raffeld M, Sharan S, Hennighausen L, Huang AM, Sterneck E., C/EBPdelta is a crucial regulator of pro-apoptotic gene expression during mammary gland involution. Development, 2005. 132(21): p. 4675–85.

38. O’Rourke, J., R. Yuan, and J. DeWille, CCAAT/enhancer-binding protein-delta (C/EBP-delta) is induced in growth-arrested mouse mammary epithelial cells. J Biol Chem, 1999. 272.

39. Hutt, J.A., J.P. O’Rourke, and J. DeWille, Signal transducer and activator of transcription 3 activates CCAAT enhancer-binding protein delta gene transcription in G0 growth-arrested mouse mammary epithelial cells and in involuting mouse mammary gland. J Biol Chem, 2000. 275.

40. Medrano, M., et al., Interrogation of Functional Cell-Surface Markers Identifies CD151 Dependency in High-Grade Serous Ovarian Cancer. Cell Rep, 2017. 18(10): p. 2343–2358.

41. Connor, P.M., et al., Characterization of the &Tumor Suppressor Pathway in Cell Lines of the National Cancer Institute Anticancer Drug Screen and Correlations with the Growth-Inhibitory Potency of 123 Anticancer Agents. Cancer Research, 1997. 57(19): p. 4285.

42. Ince, T.A., et al., Characterization of twenty-five ovarian tumour cell lines that phenocopy primary tumours. Nat Commun, 2015. 6: p. 7419.

43. Thiery, J.P., et al., Epithelial-Mesenchymal Transitions in Development and Disease. Cell, 2009. 139(5): p. 871–890.

44. Donnez, J., et al., Cyclic changes in ciliation, cell height, and mitotic activity in human tubal epithelium during reproductive life. Fertility and sterility, 1985. 43(4): p. 554–9.

45. Julie Crow, N.N.A., Jackie Lewin,Robert W. Shaw, Physiology: Morphology and ultrastructure of Fallopian tube epithelium at different stages of the menstrual cycle and menopause. Human Reproduction 1994. 9(12): p. 2224–2233.

46. George, S.H., A. Milea, and P.A. Shaw, Proliferation in the normal FTE is a hallmark of the follicular phase, not BRCA mutation status. Clinical cancer research: an official journal of the American Association for Cancer Research, 2012. 18(22): p. 6199–207.

47. Patek, E., L. Nilsson, and E. Johannisson, Scanning Electron Microscopic Study of the Human Fallopian Tube. Report II. Fetal Life, Reproductive Life, and Postmenopause**Supported by Swedish Medical Research Council Grant B72-12x-2712-04, Karolinska Institute, Stockholm, the Swedish International Development Authority, *Stockholm*, the Ford Foundation, U.S.A., and JEOL Inc., Tokyo, Japan. Fertility and Sterility, 1972. 23(10): p. 719–733.

48. Huang, A.-M., et al., The Cebpd (C/EBPd) Gene Is Induced by Luteinizing Hormones in Ovarian Theca and Interstitial Cells But Is Not Essential for Mouse Ovary Function. PLoS One, 2007(12): p. e1334–1341.

49. Davidson, B., et al., E-cadherin and α-, β-, and γ-catenin protein expression is up-regulated in ovarian carcinoma cells in serous effusions. The Journal of Pathology, 2000. 192(4): p. 460–469.

50. ElMoneim, H.M.A. and N.M. Zaghloul, Expression of E-cadherin, N-cadherin and snail and their correlation with clinicopathological variants: an immunohistochemical study of 132 invasive ductal breast carcinomas in Egypt. Clinics (Sao Paulo, Brazil), 2011. 66(10): p. 1765– 1771.

51. Davidson, B., C.G. Tropé, and R. Reich, Epithelial-mesenchymal transition in ovarian carcinoma. Frontiers in oncology, 2012. 2: p. 33–33.

52. Kalluri, R. and R.A. Weinberg, The basics of epithelial-mesenchymal transition. The Journal of clinical investigation, 2009. 119(6): p. 1420–8.

53. Trobaugh, D.W. and W.B. Klimstra, MicroRNA Regulation of RNA Virus Replication and Pathogenesis. Trends in Molecular Medicine, 2017. 23(1): p. 80–93.

54. Corney, D.C., et al., MicroRNA-34b and MicroRNA-34c Are Targets of p53 and Cooperate in Control of Cell Proliferation and Adhesion-Independent Growth. Cancer Research, 2007. 67(18): p. 8433.

55. Anandaram, H. and D.A. Anand, Computational analysis of micro RNAs compatibility in pharmacogenomic based regulatory networks of psoriatic arthritis: An initiation towards identifying a potential miRNA to treat psoriatic arthritis. Biocatalysis and Agricultural Biotechnology, 2018. 16: p. 545–547.

56. Chang, C.-J., et al., p53 regulates epithelial-mesenchymal transition and stem cell properties through modulating miRNAs. Nature Cell Biology, 2011. 13: p. 317 +.

57. Li, Y., et al., Up-regulation of miR-200 and let-7 by natural agents leads to the reversal of epithelial-mesenchymal transition in gemcitabine-resistant pancreatic cancer cells. Cancer research, 2009. 69(16): p. 6704–6712.

58. Bilyk, O., et al., Epithelial-to-Mesenchymal Transition in the Female Reproductive Tract: From Normal Functioning to Disease Pathology. Frontiers in Oncology, 2017. 7: p. 145.

59. Xu, H., et al., Let-7b-5p regulates proliferation and apoptosis in multiple myeloma by targeting IGF1R. Acta Biochim Biophys Sin (Shanghai), 2014. 46(11): p. 965–72.

60. Yu, J., et al., Let-7b inhibits cell proliferation, migration, and invasion through targeting Cthrc1 in gastric cancer. Tumour Biol, 2015. 36(5): p. 3221–9.

61. Tognon, C., T. Ng, and P.H.B. Sorensen, Reduced proliferation and enhanced migration: Two sides of the same coin? Molecular mechanisms of metastatic progression by YB-1 AU - Evdokimova, Valentina. Cell Cycle, 2009. 8(18): p. 2901–2906.

62. Tlili, S., et al., Collective cell migration without proliferation: density determines cell velocity and wave velocity. Royal Society Open Science, 2018. 5(5): p. 172421.

63. Tone, A.A., et al., Gene Expression Profiles of Luteal Phase Fallopian Tube Epithelium from BRCA Mutation Carriers Resemble High-Grade Serous Carcinoma. Clinical Cancer Research, 2008. 14(13): p. 4067–4078.

64. Ikezoe, T., et al., CCAAT/enhancer-binding protein delta: a molecular target of 1,25-dihydroxyvitamin D3 in androgen-responsive prostate cancer LNCaP cells. Cancer Res, 2005. 65.

65. Pawar, S.A., et al., C/EBP{delta} targets cyclin D1 for proteasome-mediated degradation via induction of CDC27/APC3 expression. Proc Natl Acad Sci U S A, 2010. 107(20): p. 9210–5.

66. Nik, N.N., et al., Origin and pathogenesis of pelvic (ovarian, tubal, and primary peritoneal) serous carcinoma. Annu Rev Pathol, 2014. 9: p. 27–45.

67. Vang, R., M. Shih Ie, and R.J. Kurman, Ovarian low-grade and high-grade serous carcinoma: pathogenesis, clinicopathologic and molecular biologic features, and diagnostic problems. Adv Anat Pathol, 2009. 16(5): p. 267–82.

68. Yamaguchi, J., et al., Inflammation and hypoxia linked to renal injury by CCAAT/enhancer-binding protein δ. Kidney International, 2015. 88(2): p. 262–275.

69. Kogan-Sakin, I., et al., Mutant p53(R175H) upregulates Twist1 expression and promotes epithelial–mesenchymal transition in immortalized prostate cells. Cell Death and Differentiation, 2011. 18(2): p. 271–281.

70. Wang, Y.-H., et al., CEBPD amplification and overexpression in urothelial carcinoma: a driver of tumor metastasis indicating adverse prognosis. Oncotarget, 2015. 6(31): p. 31069–31084.

71. Mittal, V., Epithelial Mesenchymal Transition in Tumor Metastasis. Annual Review of Pathology: Mechanisms of Disease, 2018. 13(1): p. 395–412.

72. Title, A.C., et al., Genetic dissection of the miR-200–Zeb1 axis reveals its importance in tumor differentiation and invasion. Nature Communications, 2018. 9(1): p. 4671.

73. Lin, S.-R., et al., MiR-193b Mediates CEBPD-Induced Cisplatin Sensitization Through Targeting ETS1 and Cyclin D1 in Human Urothelial Carcinoma Cells. Journal of Cellular Biochemistry, 2017. 118(6): p. 1563–1573.

74. May, T., et al., Low malignant potential tumors with micropapillary features are molecularly similar to low-grade serous carcinoma of the ovary. Gynecol Oncol, 2010. 117(1): p. 9–17.

75. Spandidos, A., et al., PrimerBank: a resource of human and mouse PCR primer pairs for gene expression detection and quantification. Nucleic Acids Res, 2010. 38(Database issue): p. D792– 9.

76. Wang, X. and B. Seed, A PCR primer bank for quantitative gene expression analysis. Nucleic Acids Res, 2003. 31(24): p. e154.

