## Supplementary Figures for "C/EBPδ demonstrates a dichotomous role in tumor initiation and promotion of epithelial carcinoma"

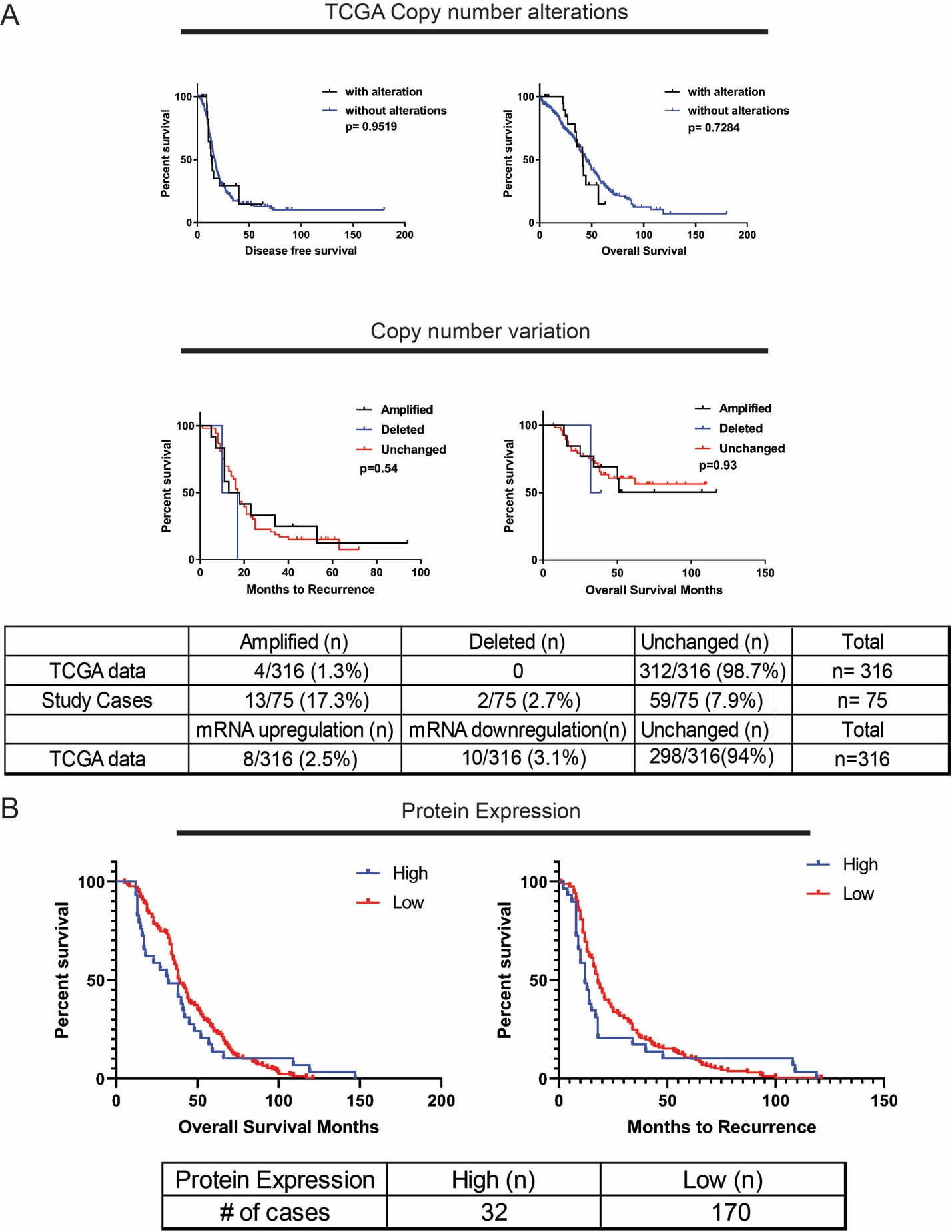

**Supplementary Figure 1. C/EBPδ copy number alteration in HGSC. A.** TCGA data showed 7% of cases with a C/EBPδ amplification or mRNA upregulation or mRNA downregulation. TCGA or our cohort of HGSC data demonstrated genetic alterations in C/EBPδ did not result in differential progression free survival or overall survival. **B.** C/EBPδ protein expression was analyzed using digital image analysis software and correlated to survival data for each patient; there are no significant differences between high and low C/EBPδ expression for overall survival (p=0.70) and month to recurrence (p=45) using the Log-rank (Mantel-Cox) test. Protein expression was divided into high (+2, moderate; +3, strong) and low (+1, weak).

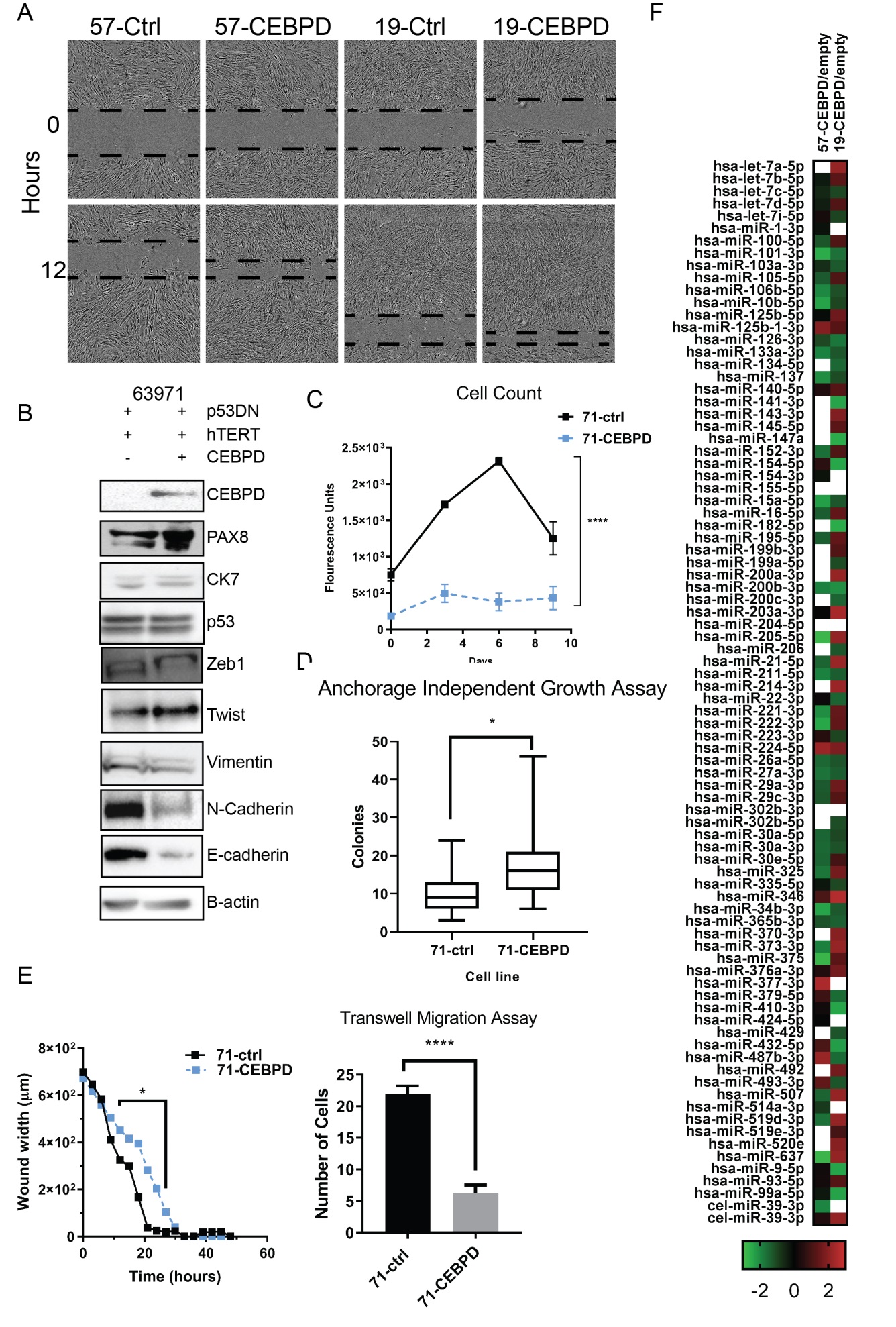

**Supplementary Figure 2.** **A**. The migratory potential of FTE-57 and FTE-19 was determined using a wound healing assay. Cells with CEBPD-OE migrated quicker relative to control cells (57-p53DN-hTert (2.04-fold, p<0.01) and 19-p53DN-hTert (1.6-fold, p<0.01)). **B.** Western blot analysis of EMT markers FTE-63971. **C.** Over-expression of C/EBPδ also decreased cell growth of FTE-63971 similar to other FTE cell lines (Statistical significance derived using paired t-test (p<0.0001)). **D.** Increase in colony formation in this more mesenchymal phenotype FTE-line when C/EBPδ is over-expressed. **E.** An inverse in migration is seen when E-cadherin is not up-regulated – a decrease in both migration and invasion when C/EBPδ is over-expressed **F.** miRNA expression between FTE with C/EBPδ and without C/EBPδ was assessed and showed many miRNA differentially expressed between FTE-CEBPD and FTE-control cells. Statistical significance derived using 2-way ANOVA. Statistically significant (p<0.05) miRNA placed in a shortlist and p-values are determined by a Student’s t-test (two tail distribution and equal variances between the two samples).

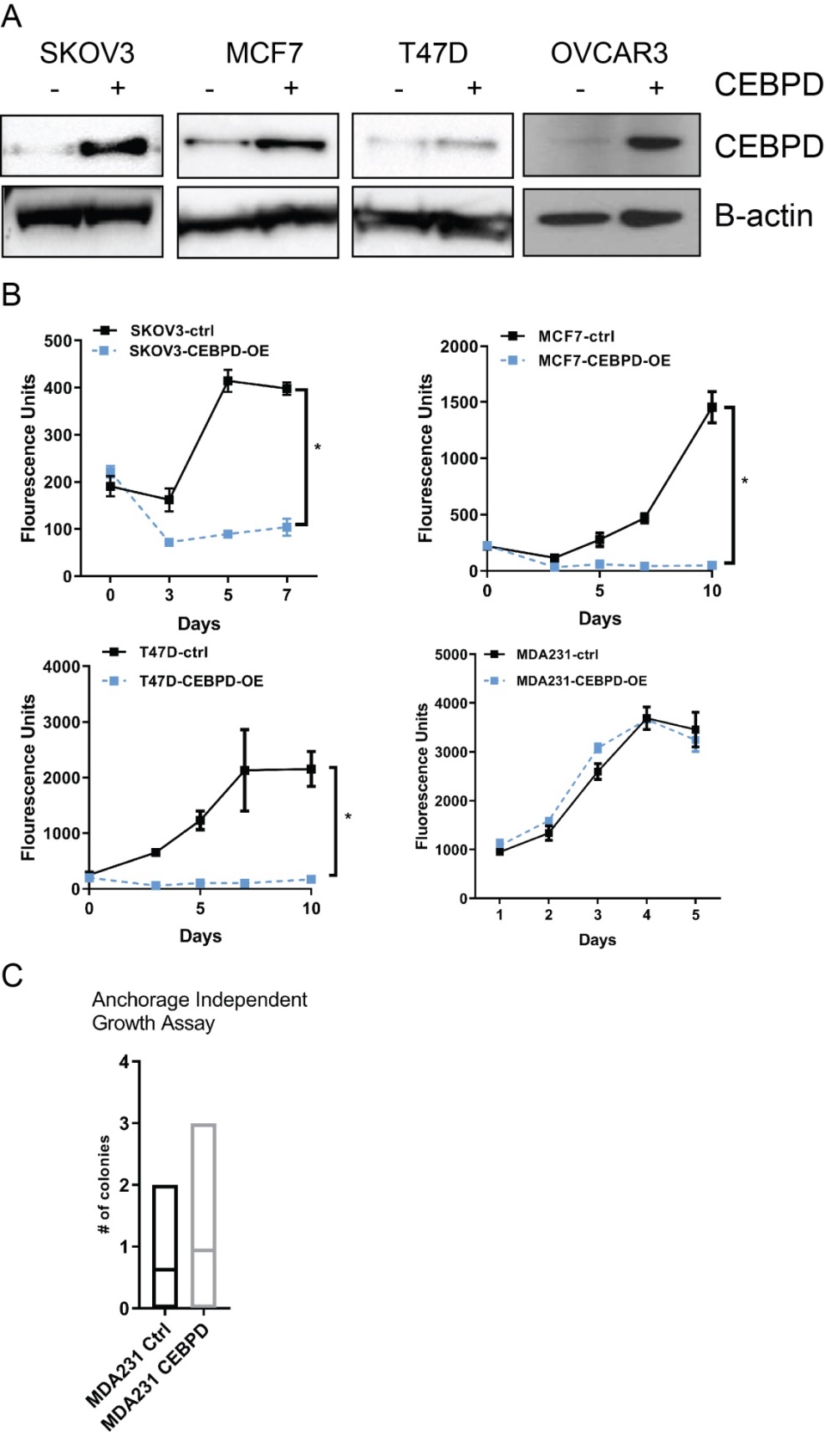

**Supplementary Figure 3.A.** A western blot of SKOV3, MCF7,T47D and OVCAR3 overexpressed with C/EBPδ confirming protein expression. **B.** Proliferation assays performed on several cancer cell lines to assess cell growth. **C.** A soft agar assay performed on MDA231 cells and the number of colonies between CEBPD-OE and control cells. Data are represented with statistical signifcance set at p< 0.05.

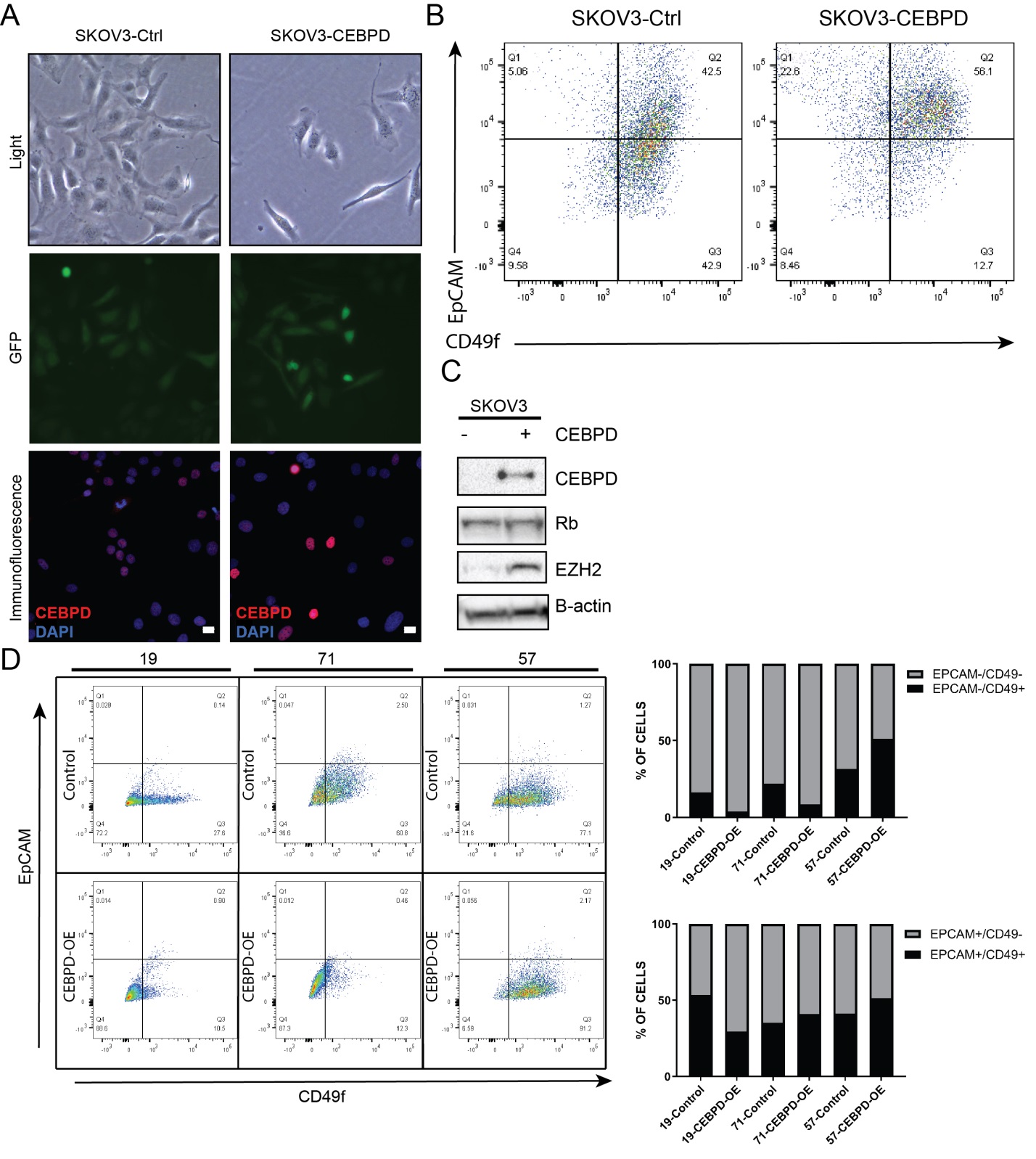

**Supplementary Figure 4. A.** C/EBPδ expressed in SKOV3 cells and confirmed by GFP and immunofluorescence. **B.** FACS analysis of SKOV3 cells show CEBPD-OE cells with increased EPCAM and CD49f expression relative to control cells. **C.** Overexpression of C/EBPδ did not alter Rb levels, but increased EZH2 relative to control cells. **D.** FACS analysis of three fallopian cells show EPCAM and CD49 expression in cells expressing C/EBPδ vs control.

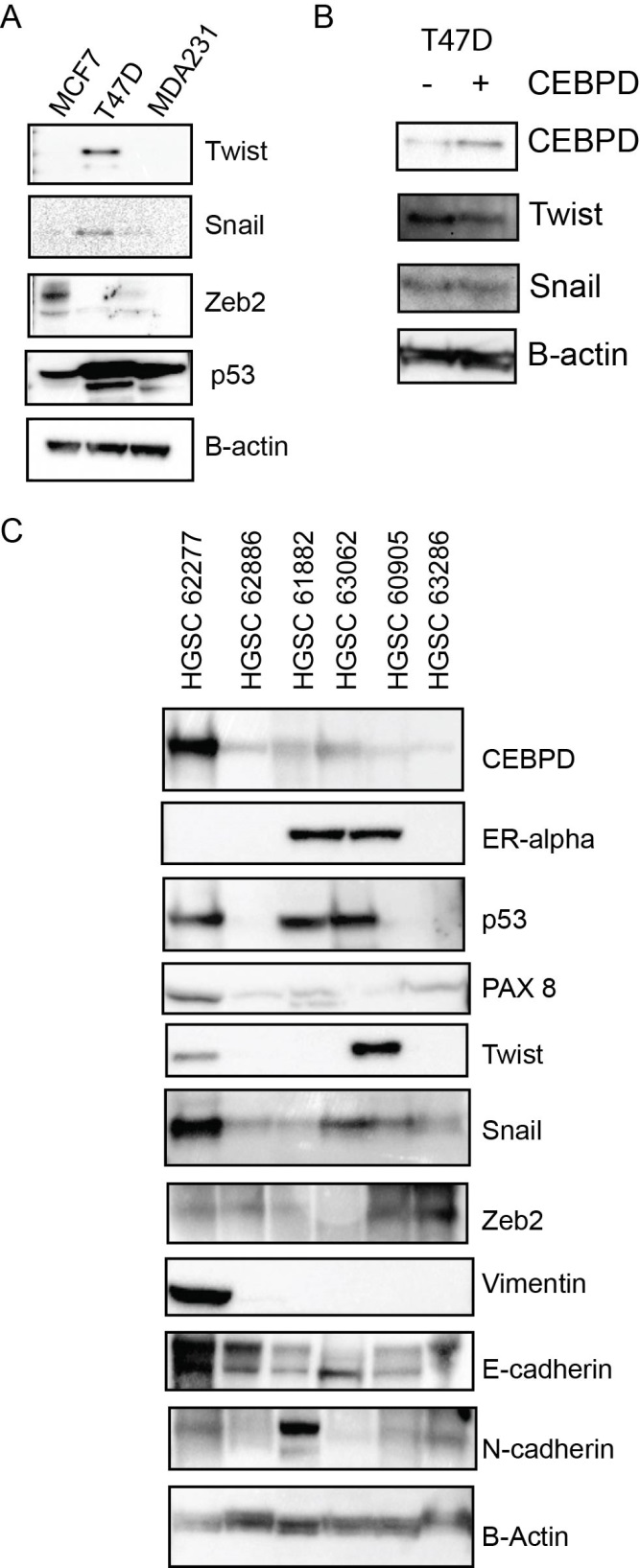

**Supplementary Figure 5.** **A.** Western Blot Images of three breast cancer cell lines (MCF7, T47D, MDA-231) show differential expression of Twist, Snail, Zeb2, p53 and B-actin. **B.** Additional breast cancer cell lines show western blot protein expression level of C/EBPδ and EMT markers. **C.** Western blot of six HGSC and expression levels of EMT/MET markers.

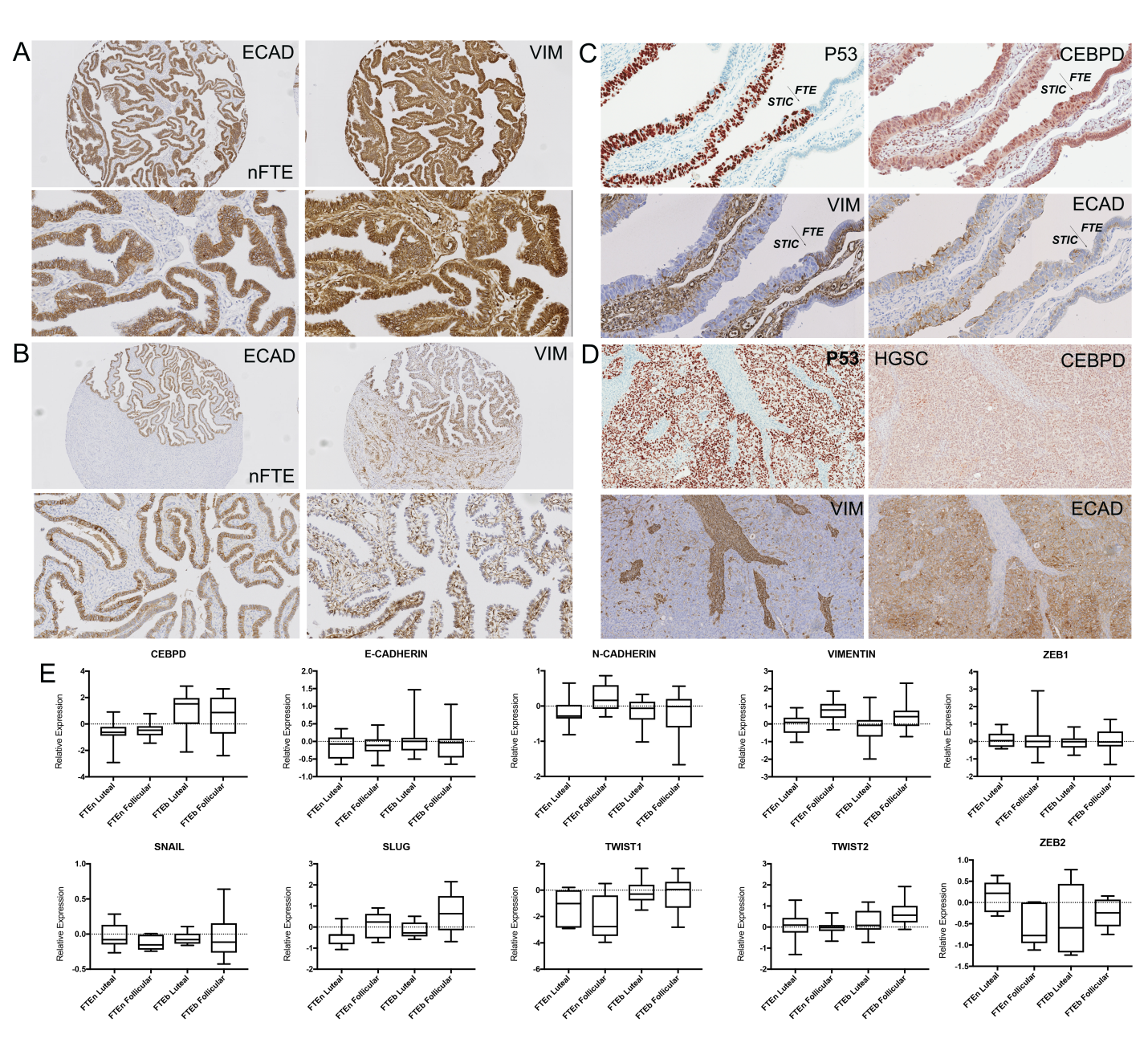

**Supplementary Figure 6.** **A.** High expression level of E-cadherin (ECAD) and Vimentin (VIM) in normal fallopian tube epithelium. **B.** Low expression level of E-cadherin and Vimentin in normal fallopian tube epithelium. **C.** Comparative immunohistochemical image of C/EBPδ, E-cadherin and Vimentin, showing differential expression of each marker between normal fallopian tube epithelia and serous tubal intraepithelial carcinoma. **D.** Expression level of C/EBPδ, E-cadherin and Vimentin in HGSC. **E.** Normalized gene expression data derived from microarray analysis of normal fallopian tube cases of the luteal and follicular phases and BRCA mutation carriers of the luteal and follicular phases.
